# Unfolding spatiotemporal representations of 3D visual perception in the human brain

**DOI:** 10.1101/2025.08.03.668371

**Authors:** Zitong Lu, Julie D. Golomb

## Abstract

Although visual input is initially recorded in two dimensions on our retinas, we perceive and interact with the world in three dimensions. Achieving 3D perception requires the brain to integrate 2D spatial representations with multiple depth cues, such as binocular disparity. However, most studies typically examine 2D and depth information in isolation, leaving the integrated nature of 3D spatial encoding largely underexplored. In this study, we collected a densely sampled multimodal neuroimaging dataset from 10 participants (8 with EEG and fMRI; 2 with fMRI only) across multiple sessions while they viewed stereoscopic 3D stimuli through red-green anaglyph glasses. Participants first completed a behavioral session including depth judgement tasks and a novel cube adjustment task to quantify and calibrate individual depth perception in units of binocular disparity. Then during two EEG and two fMRI sessions, participants passively viewed stimuli presented at 64 systematically sampled 3D locations, yielding over 66,000 trials in total across ten participants. Combining this multimodal dataset with computational methods via representational similarity analysis, we examined how 2D, depth-related, 3D feature-level, and geometric distance representations unfold across time and brain space. We found that the human brain represents 3D visual space not only by encoding position-in-depth as an additional dimension alongside 2D location, but also by constructing richer forms of 3D spatial structure. Specifically, 2D spatial features were represented earliest and most broadly, depth-related representations were weaker and more spatially restricted, and 3D feature representations were sparse and heterogeneous but detectable at the individual feature level. Critically, geometric distance analyses revealed that neural coding extended beyond separable feature dimensions, showing robust integrated 2D geometric representations and more selective evidence for integrated 3D geometric representations. These findings suggest that human 3D spatial perception is supported by a progression from dominant 2D coding to depth-related and 3D representations, with additional evidence for geometric structure in 3D space, contributing to a more comprehensive understanding of the spatiotemporal organization of neural representations that support 3D perception. Additionally, our novel large dataset will be made openly available to support future research on 3D perception and spatial cognition.

## Introduction

Despite the inherently two-dimensional nature of retinal input, humans experience the world as a richly structured three-dimensional environment. The human visual system integrates 2D spatial representations with a range of depth cues– including binocular disparity, perspective, shading, relative size, occlusion, and motion – to construct rich 3D spatial percepts that support perception, navigation, and interaction with the environment (Howard, 2012; Welchman, 2016). Yet an important conceptual question remains unresolved: does the human brain represent 3D space merely by encoding position-in-depth as an additional spatial dimension alongside 2D location, or does it also construct genuinely higher-order 3D representations that go beyond these component dimensions? Resolving this central question is essential for understanding how the visual system moves from a 2D retinal image to a behaviorally useful representation of the 3D world.

Decades of research have investigated how the brain encodes 2D spatial dimensions of space (T. Carlson et al., 2011; Engel et al., 1994; Fischer et al., 2011; Golomb & Kanwisher, 2012; Grill-Spector & Malach, 2004; Kravitz et al., 2010; Maunsell & Newsome, 1987; Schwarzlose et al., 2008; Sereno et al., 1995; Silver & Kastner, 2009; Tootell et al., 1998; Wandell et al., 2007), as well as how visual areas respond to depth information conveyed by binocular disparity and other cues (Backus et al., 2001; Bridge et al., 2023; Chen et al., 2020; Ip et al., 2014; Neri et al., 2004; Uka & DeAngelis, 2006). Recent findings further suggest a gradual transition from 2D to depth representations along the visual hierarchy (Finlayson et al., 2017; Henderson et al., 2019). However, most prior work has investigated or analyzed 2D and depth in isolation, leaving unclear whether the human brain represents 3D space only through depth as an additional dimension, or whether it also forms richer 3D representations that integrate these dimensions into 3D-related and more complicated spatial codes.

A major obstacle to addressing this question is that perceived depth cannot be directly inferred or measured from binocular disparity in a uniform way across observers. Fixed disparity values do not necessarily correspond to the same perceived depth magnitude across individuals, making it difficult to place depth on the same metric scale as horizontal and vertical position. As a result, previous studies using fixed disparity levels have had limited ability to test whether neural responses reflect depth as an additional dimension or richer forms of 3D spatial representation (Alvarez et al., 2021; Bridge & Parker, 2007; Finlayson et al., 2017). In addition, resolving this issue requires dense sampling of locations across a calibrated 3D space, together with neural measurements that can reveal both where and when different representational formats emerge.

Here, we overcame these challenges using a multimodal framework for tracking neural representations of calibrated 3D space across time and brain space. To precisely quantify perceived depth from binocular disparity, participants first performed a behavioral “cube adjustment” task in which participants dynamically adjusted the distances between 3D stimuli to match perceived horizontal, vertical, and depth distances; i.e., until the stimuli formed the corners of a perfect cube (Figure 1A). This individualized calibration allowed us to derive a parametric mapping between physical disparity and perceived 3D depth for each participant, such that we could construct a participant-specific 3D stimulus space in which depth could be placed on a common perceptual scale with horizontal and vertical position. We then collected an extensive dataset of 3D visual perception comprising both EEG and fMRI from multiple sessions per participant (Figure 1A). Stimuli for these neuroimaging sessions were viewed stereoscopically through red-green anaglyph glasses and sampled a structured 3D space with 64 distinct spatial locations (4×4×4 grid centered on fixation). By integrating individualized depth calibration, densely sampled multimodal EEG and fMRI recordings, and representational similarity analysis (RSA) (Figure 1B-E; also see more details in Methods), we tested whether neural representations of 3D space are limited to 2D location plus depth as component feature dimensions, or whether the human brain also encodes more higher-order forms of 3D spatial structure. As additional exploratory questions, we further examined whether these representations show preferences for different spatial coordinate formats across time and brain space, and how individual differences in depth perception relate to neural 3D spatial coding. Together, this multimodal, publicly available dataset provides a framework for characterizing how 2D, depth, and more integrated 3D representations unfold across time and brain space.

**Figure 1.**
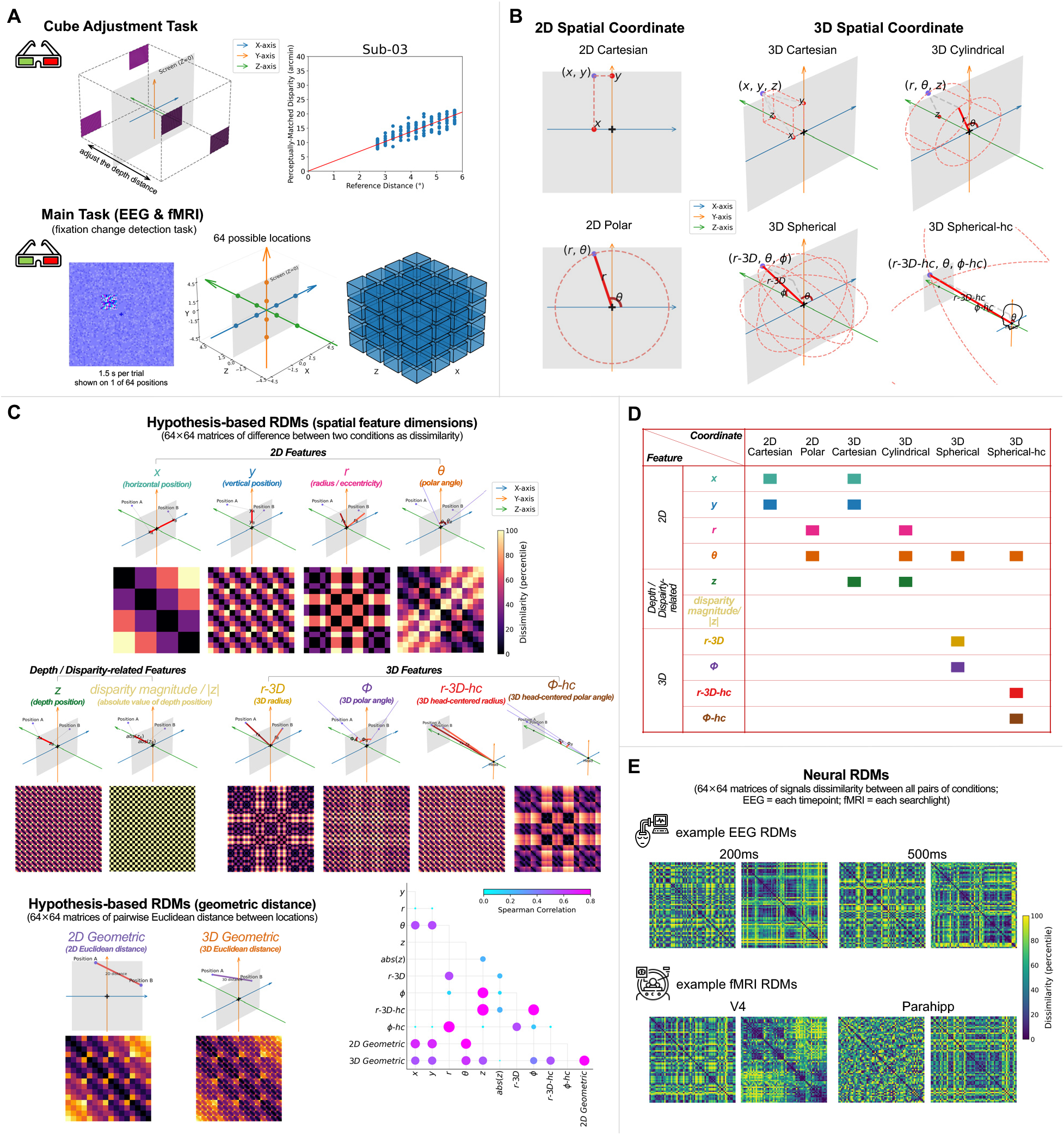
Experimental design, spatial coordinate systems, and representational patterns. (A) Tasks in our study. In the Cube Adjustment Task for individualized depth calibration, participants adjusted the depth distance between the front and back panels using the vertices of the stimuli to generate a virtual cube, allowing us to derive a personalized mapping between binocular disparity and perceived depth distance. Upper-left: Schematic of the cube adjustment task. Upper-right: Example calibration function fitted based on one subject’s (Sub-03) real behavioral data. In the Main EEG and fMRI task, participants performed a fixation-change detection task while passively viewing stimuli presented at one of 64 3D spatial locations arranged in a 4×4×4 grid. Stimuli were small cubes of high-contrast dynamic random dot stimuli presented on a lower-contrast static background viewed through red-green anaglyph glasses. (B) Schematics illustrating the possible 2D and 3D spatial coordinate systems used to define spatial features. For each schematic, the black fixation cross indicates the fixation location and the gray dot the example stimulus location, with the light gray rectangle depicting the computer monitor. (C) Hypothesis-based RDMs (64 × 64 matrices). Ten spatial feature RDMs were constructed for each of the ten different spatial feature dimensions by calculating pairwise differences in feature values between conditions. Two geometric distance RDMs were constructed by calculating pairwise Euclidean distances among locations in either the 2D frontoparallel plane or the full calibrated 3D space. Top schematics show geometric representations of an example pair of stimuli (Position A and Position B) and how dissimilarity is calculated for each hypothesis (red lines; e.g. comparison of *x*_A_ and *x*_B_). Bottom schematics show the 64×64 RDMs for each corresponding hypothesis. Spearman correlations between these hypothesis-based RDMs are shown in the inset on the bottom right. (D) Table showing the spatial coordinate systems (columns) and which spatial feature dimensions (rows) they contain. Colored squares indicate the constituent features of each 2D or 3D spatial coordinate. (E) Example Neural RDMs. Neural RDMs were computed separately for each EEG time point or each fMRI searchlight, capturing pairwise signal dissimilarity between the 64 spatial conditions. For each example, both amplitude-based (left) and pattern-based RDMs (right) are shown. Example EEG RDMs are shown at selected time points (200ms and 500ms), and example fMRI RDMs are shown from selected cortical regions (V4 and parahippocampus).

## Results

### Analytical framework for probing human 3D spatial representations

To investigate how the human brain represents 3D visual space, we first established an individualized mapping between binocular disparity and perceived depth for each participant using the behavioral cube-adjustment task (Figure 1A). In this task, participants adjusted the front-back separation of stereoscopic stimuli until the four stimulus vertices were perceived as forming corners of a cube, allowing us to estimate the disparity required to produce matched perceived depth distances for each participant (see each participant’s results in Figure S1). We then used this participant-specific calibration to construct 64 stimulus locations in the main EEG and fMRI experiments, such that depth could be placed on a common perceptual scale with horizontal and vertical position. This design allowed us to examine neural representations of spatial location in a perceptually calibrated 3D space, rather than relying on fixed physical disparity values alone.

We applied representational similarity analysis (RSA) to test whether neural representations of 3D space are limited to separable 2D and depth-related dimensions, or whether the human brain also encodes higher-order forms of 3D spatial structure. We constructed hypothesis-based representational dissimilarity matrices (RDMs) capturing individual spatial features, including horizontal position (*x*), vertical position (*y*), depth position (*z*), radius/eccentricity (*r*), polar angle (*θ*), 3D radius (*r-3D*), 3D polar angle (*Φ*), 3D head-centered radius (*r-3D-hc*), and 3D head-centered polar angle (*Φ-hc*), derived from six possible spatial coordinate systems (2D Cartesian, 2D Polar, 3D Cartesian, 3D Cylindrical, 3D Spherical, and 3D Spherical-hc) across the 64 spatial locations (Figure 1B-D). To distinguish perceptual depth-related coding from effects driven by absolute disparity magnitude, we also included a binocular disparity magnitude (|*z*|) RDM based on the absolute value of depth position. We also constructed two RDMs reflecting hypothetical integrated forms of spatial structure: a 2D geometric-distance RDM and 3D geometric distance RDM, based on pairwise Euclidean distances among locations in either the 2D frontoparallel plane or the full calibrated 3D space (Figure 1C), to test whether neural representations reflect spatial relationships beyond separable feature dimensions.

We computed neural RDMs from the EEG data at each time point and from the fMRI data at each searchlight unit or ROI (Figure 1E), and then computed partial correlations between neural RDMs and each hypothesis-based RDM. For the spatial feature analyses, this allowed us to estimate the unique contribution of each spatial feature after accounting for shared variance with other feature dimensions. For the geometric-distance analyses, this allowed us to test whether neural activity reflects integrated 2D or 3D spatial relationships, including whether these effects persisted after accounting for component feature-level coding. Here, RSA was used not to identify isolated spatial parameters per se, but to test structured hypotheses about the organization of spatial representations. Specifically, by comparing neural RDMs with hypothesis-based model RDMs, we assessed whether spatial information is represented as separable component dimensions (e.g., 2D position and depth) or as more integrated forms of spatial structure. We first asked whether neural activity at different points of time and brain locations contains distinct representations of 2D, depth, and 3D spatial features. Then we tested whether spatial coding extends beyond these separable component dimensions to more integrated geometric representations, and finally treated coordinate-format preferences, brain-behavior relationships, and EEG-fMRI representational correlations as exploratory analyses.

For both EEG and fMRI, we constructed two complementary types of neural RDMs: amplitude-based RDMs and pattern-based RDMs. Amplitude-based RDMs capture differences in overall response magnitude across conditions, whereas pattern-based RDMs capture differences in multivariate spatial activity patterns across channels or voxels. Because these two RDM types reflect distinct aspects of neural coding, we analyzed both separately. In the main figures, significant EEG time points are marked according to which RDM type primarily drove the effect: open circles indicate effects driven primarily by amplitude-based RDMs, whereas filled circles indicate effects driven primarily by pattern-based RDMs. For fMRI ROI analyses, light-gray asterisks indicate effects driven primarily by amplitude-based RDMs, whereas dark-gray asterisks indicate effects driven primarily by pattern-based RDMs. Here, “primarily driven by” refers to the RDM type with the larger effect among the two, and does not imply that the other RDM type showed no effect. Full results for amplitude-based and pattern-based RDMs are reported separately in the Supplementary Figures.

### A spatiotemporal hierarchy from 2D to depth and 3D spatial representations

We first asked, at a broad representational level, about the nature of neural coding beyond 2D space. Prior studies have revealed a cortical hierarchy where representations appear to transition from robust 2D representations to representations containing some depth information (e.g., Finlayson et al., 2017) – a key question is whether neural coding beyond 2D space is limited to depth position alone or whether it also contains richer 3D spatial information that depends on the joint organization of horizontal, vertical, and depth positions, and cannot be reduced to any single 2D or depth dimension. To address this question, as illustrated in Figure 1C, our hypothesis-based spatial feature RDMs were organized into three categories: 2D spatial features (*x*, *y*, *r*, *θ*), depth/disparity-related features, and 3D spatial features. The depth/disparity-related category included signed position-in-depth (z) together with an absolute disparity-magnitude control model (|z|), which was included to account for effects driven by disparity magnitude rather than being interpreted on its own as perceived depth coding. The 3D category included spatial features that are defined only in a 3D coordinate space and not reducible to any single 2D or depth dimension (*r-3D*, *ϕ*, *r-3D-hc*, *ϕ-hc*).

Because different features within the same category could emerge at different times or in different brain regions, averaging across features might obscure sparse or heterogeneous effects. We therefore used category-wise maximum and presence analyses to ask whether any feature within each category was represented at a given time point or cortical location. For EEG, we extracted the maximum representational similarity across features within each category at each time point. For fMRI, we identified cortical locations where at least one feature within each category was significantly represented.

The EEG results revealed a clear temporal hierarchy across feature categories (Figure 2A). 2D features showed the strongest and earliest representational similarity, emerging rapidly after stimulus onset and remaining significant across an extended time window. In contrast, depth/disparity-related features showed weaker and more temporally restricted effects. Importantly, 3D features were also detectable, despite being substantially weaker than the dominant 2D response. These effects emerged later and were more temporally specific, suggesting that neural coding beyond the frontoparallel plane is present but less robust than 2D spatial coding. To quantify this temporal progression, we compared onset and peak latencies across the three feature categories (Figure 2B). Consistent with the descriptive time-course results, 2D features showed the earliest onset and peak latencies. Depth/disparity-related and 3D features generally showed later peak latencies, with 3D features peaking latest overall. This pattern suggests a progression from rapid and sustained 2D spatial coding toward weaker and later coding of depth/disparity-related and 3D spatial information.

**Figure 2.**
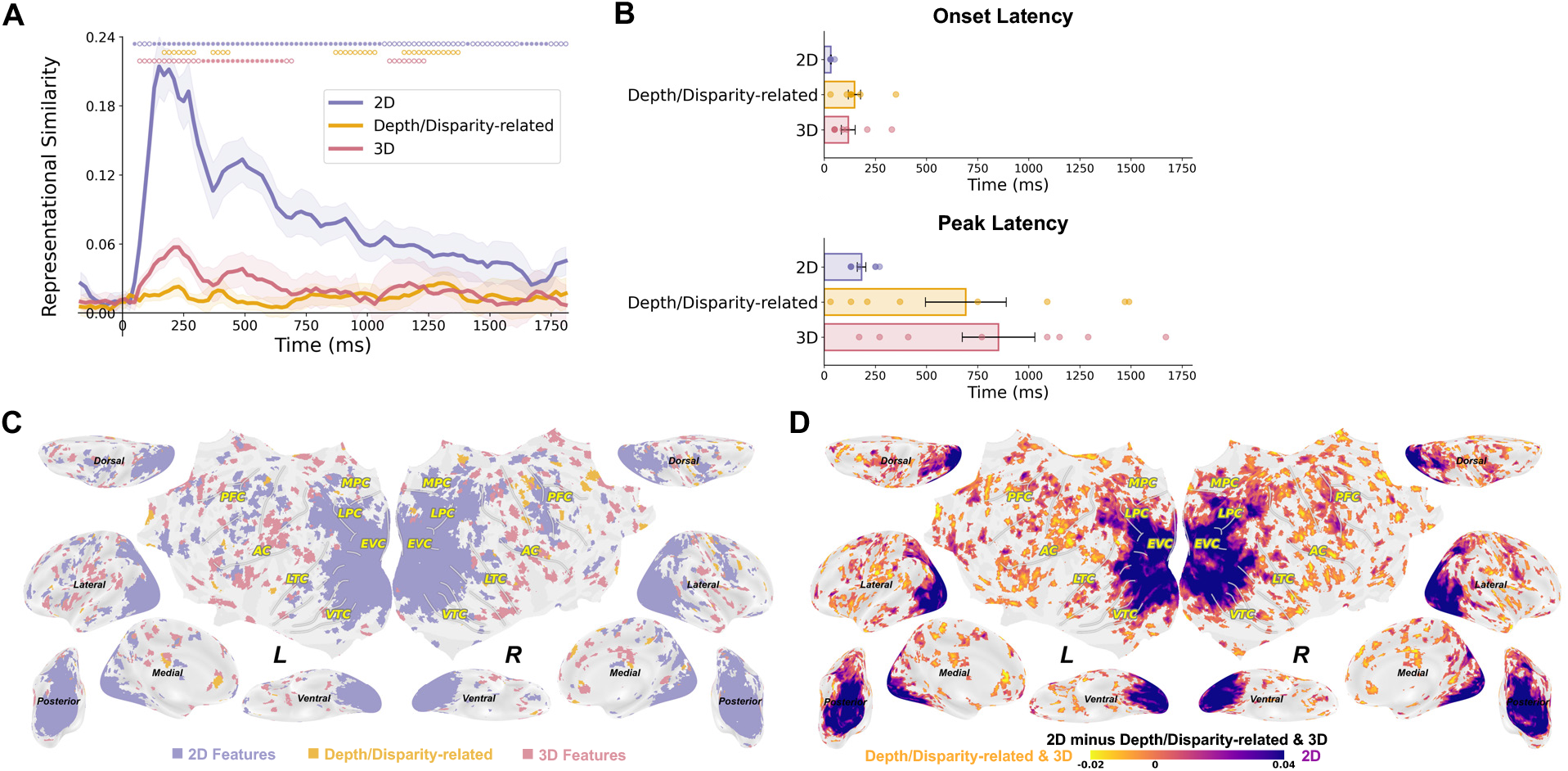
Category-wise maximum and presence analyses of 2D, depth/disparity-related, and 3D feature representations. (A) EEG time sources of the maximum representational similarity (partial Spearman correlation) across features within each category: 2D features (*x*, *y*, *r*, *θ*), depth/disparity-related features (*z* and |*z*|), and 3D features (*r-3D*, *ϕ*, *r-3D-hc*, *ϕ-hc*). Shaded areas indicate ±1 SEM across participants. Open circles indicate significant time points primarily driven by amplitude-based RDMs, whereas filled circles indicate significant time points primarily driven by pattern-based RDMs (permutation tests with correction for the number of hypothesis-based RDMs, followed by cluster-based correction, *p*<.05). (B) Onset and peak latencies of the category-wise maximum EEG representational similarity (partial Spearman correlation) results. Dots denote individual participants; error bars indicate SEM. (C) fMRI category-presence map showing cortical locations where at least one feature within a category was significant (permutation tests with correction for the number of hypothesis-based RDMs, *p*<.05). Each cortical location is labeled according to the presence of significant 2D, depth/disparity-related, or 3D feature representations. (D) Direct contrast between the strongest 2D feature representation and the strongest higher-dimensional representation (depth/disparity-related and 3D features) at each voxel. Positive values indicate stronger 2D representations, whereas negative values indicate stronger depth/3D representations.

The fMRI searchlight results showed a complementary spatial hierarchy. We first generated a category-presence map in which a cortical location was labeled as 2D, depth/disparity-related, or 3D if at least one feature within that category was significant at that location (Figure 2C). This analysis showed widespread 2D feature representations and more sparse, distributed depth/disparity-related and 3D feature representations. To further characterize this spatial organization, we then directly contrasted the strongest 2D feature representation against the strongest higher-dimensional representation at each voxel, where higher-dimensional representations were defined as the maximum across both depth/disparity-related and 3D features (Figure 2D; see also Figure S2). By incorporating a broader set of spatial features, this contrast suggested a spatial gradient in feature-level coding: early visual cortex and adjacent posterior visual regions were dominated by 2D representations, whereas more anterior and higher-order cortical regions showed relatively stronger depth/disparity-related and 3D representations.

This spatiotemporal difference was also evident in the feature-count analysis (Figure S3). Whereas the widespread 2D feature representations were supported by multiple 2D features across broad posterior visual regions, 3D feature representations were more heterogeneous: different cortical areas appeared to be driven by different 3D features, with relatively few regions showing convergent evidence across multiple 3D dimensions. Thus, the 3D representation map should not be interpreted as reflecting a single uniform 3D representation across cortex, but rather as revealing sparse and spatially heterogeneous 3D feature coding.

Together, these results reveal a broad spatiotemporal hierarchy in the neural representation of 3D visual space. 2D spatial features are represented earliest, most strongly, and most broadly across cortex. Depth/disparity-related and 3D features are weaker, later, and more spatially restricted, but they are not absent. This pattern suggests that human 3D spatial perception is supported by dominant frontoparallel spatial coding together with more selective coding of depth/disparity-related and 3D-derived spatial features. In the next section, we examine which specific spatial features drive these effects, before turning to our other primary theoretical question: whether spatial coding extends beyond these separable spatial feature component dimensions to more integrated geometric representations.

### Spatiotemporal coding at the level of individual spatial features

Having established the broad category-level hierarchy, we next examined which individual spatial features contributed to these effects across time and brain space. Because these analyses were based on partial correlations, each feature-level result reflects representational variance that was uniquely associated with that feature after removing variance shared with the other hypothesis-based spatial feature RDMs.

Several feature-specific patterns were especially prominent (Figures 3 and 4; Combined visualization of EEG and fMRI results are also illustrated in Supplementary Video 1). Among the 2D features, polar angle (*θ*) and radius/eccentricity (*r*) showed the strongest and earliest EEG effects, emerging shortly after stimulus onset and persisting across much of the epoch. These temporal effects were accompanied by broad fMRI representations across posterior visual cortex and higher visual regions, consistent with robust and spatially widespread coding of these 2D spatial dimensions. Horizontal and vertical position also showed significant representations, but these effects were generally weaker and more spatially restricted than *θ* and *r*.

**Figure 3.**
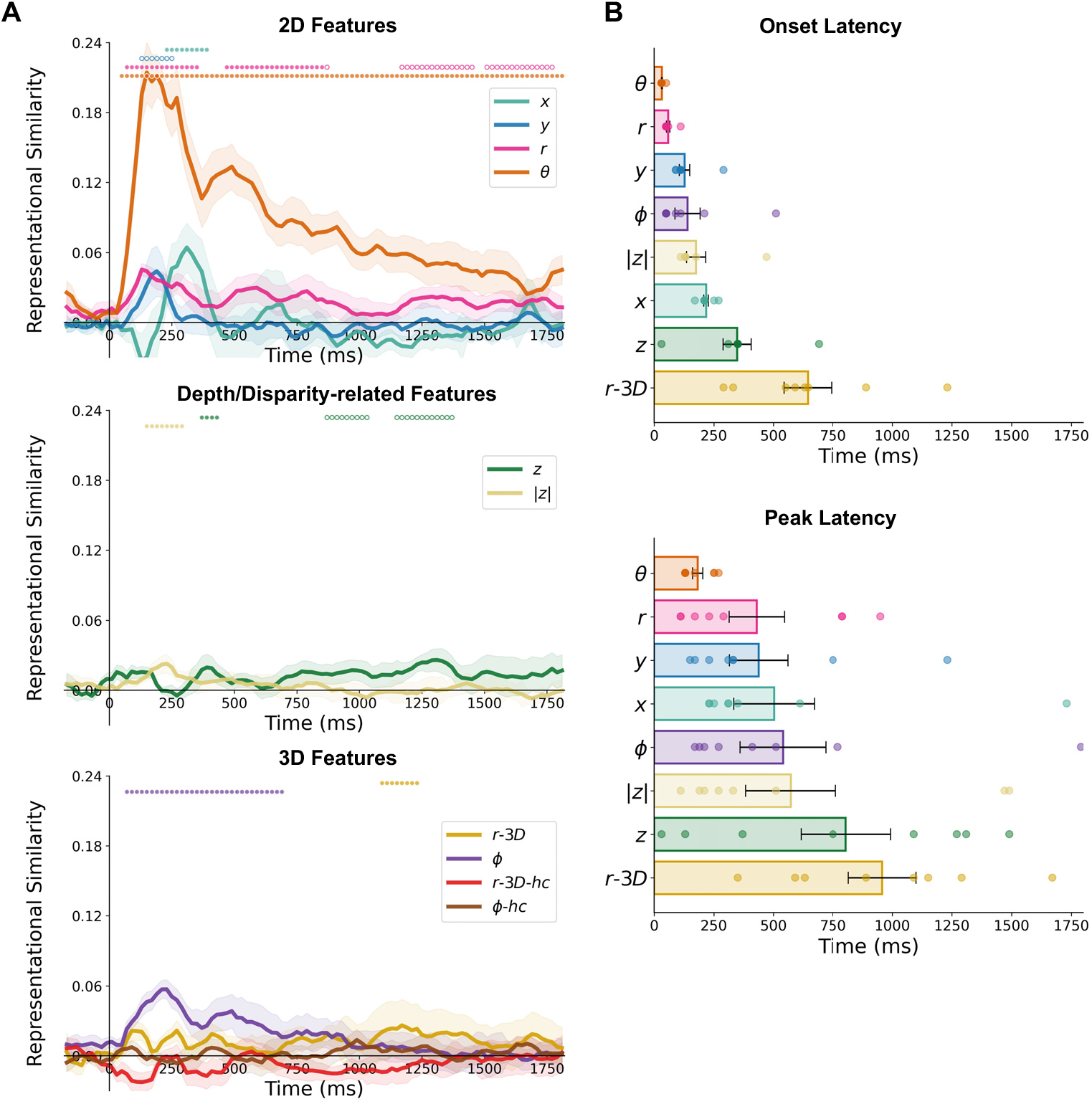
Feature-level temporal dynamics of 2D, depth/disparity-related, and 3D spatial representations. (A) Time-resolved representational similarity (partial Spearman correlation) between EEG temporal RDMs and the ten hypothesis-based spatial feature RDMs, including 2D features (*x*, *y*, *r*, *θ*), depth/disparity-related features (*z* and |*z*|), and 3D features (*r-3D*, *ϕ*, *r-3D-hc*, *ϕ-hc*). Shaded areas indicate ±1 SEM across participants. Open circles indicate significant time points primarily driven by amplitude-based RDMs, whereas filled circles indicate significant time points primarily driven by pattern-based RDMs (permutation tests with correction for the number of hypothesis-based RDMs, followed by cluster-based correction, *p*<.05). (B) Onset and peak latencies of EEG representational similarity (partial Spearman correlation) effects for spatial features that showed significant time points. Features without significant EEG effects (*r-3D-hc* and *ϕ-hc*) were not included in the latency summary. Dots denote individual participants; error bars indicate SEM.

**Figure 4.**
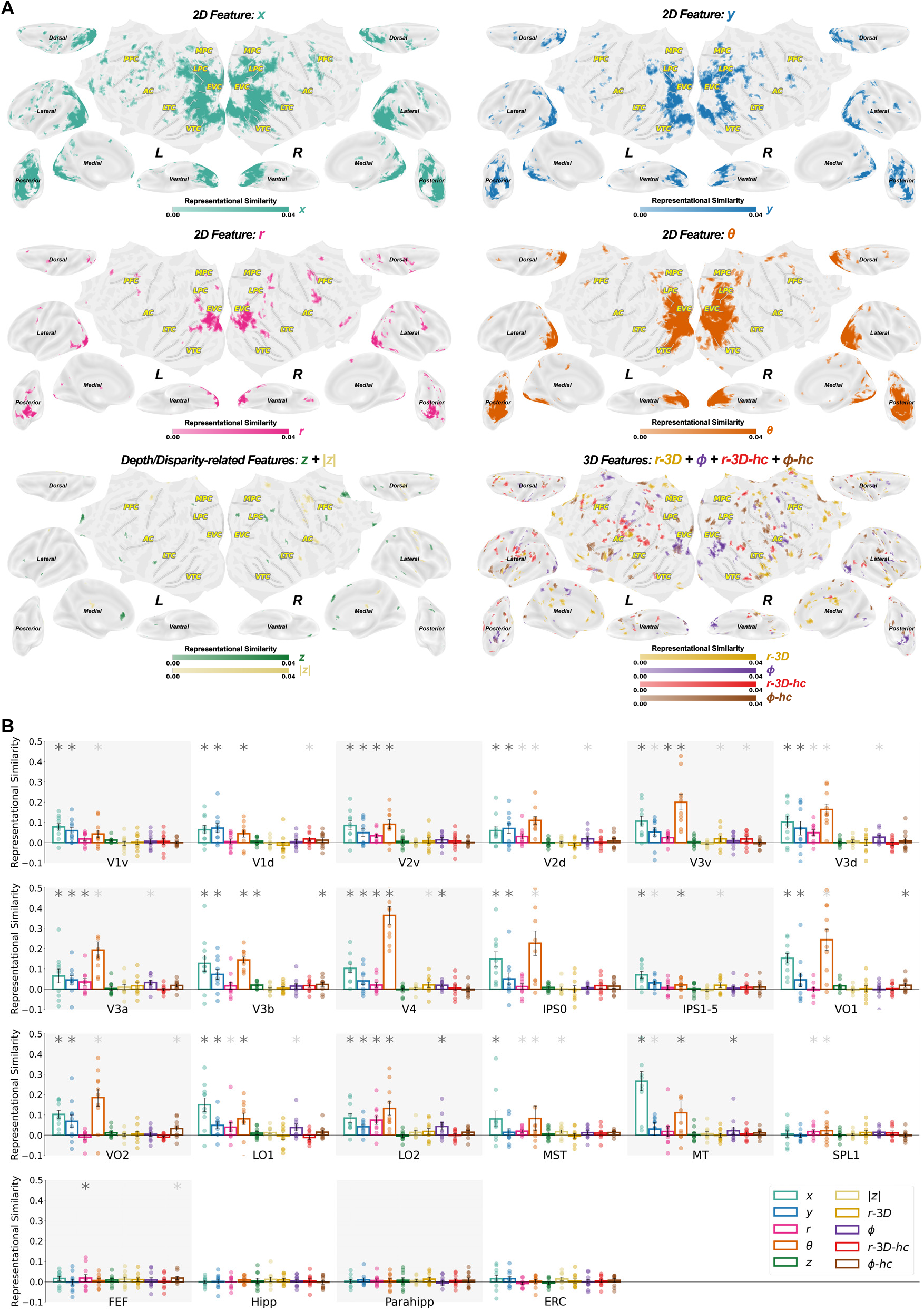
Spatial distribution of feature-level representations across brain space. (A) fMRI searchlight representational similarity (partial Spearman correlation) results for spatial features displayed on inflated cortical surfaces. The four 2D feature maps (*x*, *y*, *r*, *θ*) are shown separately, whereas depth/disparity-related features (*z* and |*z*|), and 3D features (*r-3D*, *ϕ*, *r-3D-hc*, *ϕ-hc*) are summarized as two combined presence maps because their individual maps were more sparse and heterogeneous. Colored voxels indicate the searchlight centered on that voxel was significant (permutation tests with correction for the number of hypothesis-based RDMs, followed by cluster-based correction, *p*<.05). (B) ROI-based RSA results showing group-level representational similarity (partial Spearman correlation) across 25 predefined ROIs. Light-gray asterisks indicate significant effects primarily driven by amplitude-based RDMs, whereas dark-gray asterisks indicate significant effects primarily driven by pattern-based RDMs (permutation tests with correction for the number of hypothesis-based RDMs, *p*<.05). Dots denote individual participants. Error bars indicate SEM.

Depth/disparity-related and 3D features showed more selective feature-specific profiles. *z* and |*z*| showed weaker but temporally extended EEG effects, with more restricted fMRI representations than the dominant 2D features. Among the 3D features, *ϕ* and *r-3D* showed significant EEG effects, with *ϕ* emerging in an early-to-mid time window and *r-3D* showing a more temporally specific profile. In fMRI, different 3D features were represented in partially distinct cortical locations: for example, *ϕ* was more evident in occipital and visual cortical regions, whereas other 3D features such as *r-3D*, *r-3D-hc*, and *ϕ-hc* appeared more sparsely and in more distributed regions. Thus, the individual feature-level results suggest that 3D feature coding is not expressed as a single uniform 3D feature map, but rather as a heterogeneous set of partially distinct feature representations across time and brain space.

Taken together, these feature-level analyses enhance the initial results by clarifying which specific spatial dimensions underlie the broader category-level effects described above. Because these results were obtained using partial-correlation RSA, they reflect unique feature contributions after accounting for shared variance among spatial models. For comparison, the corresponding correlation-based RSA results, in which shared variance was not removed, are shown in Supplementary Figure S4–S5.

### Geometric-distance representations of 2D and 3D space

While the preceding analyses focused on the encoding of individual spatial features, it remains unclear whether the brain also processes spatial location in an integrated, geometric manner – that is, whether it could form a unified 2D or 3D positional coding pattern which reflects not the individual feature dimensions but their combined geometric relationships in space. For example, two objects that differ by one unit horizontally and one unit vertically would be separated by √2 units in an integrated 2D Euclidean space. A feature-based encoding scheme might treat these two objects as distinct along independent horizontal and vertical maps, while a geometric distance representation would capture their spatial dissimilarity as a function of their combined distance in this space: how similar or dissimilar two locations are in 2D or 3D space. RSA is well suited for testing this question because it directly asks whether the pairwise dissimilarity structure of neural activity tracks the pairwise geometric structure of the stimulus space, and can reveal time points and brain regions in which neural coding scheme is more reflective of feature-based neural encoding or geometric distance representations. Critically, such geometric distance representations highlight a fundamentally different kind of spatial information. While feature-based maps may support tasks like localizing and reaching toward a single object, geometric-distance-based codes may underlie the perception of global spatial structure and inter-object relationships.

To test this, we constructed two geometric distance RDMs (Figure 2C): a 2D geometric distance RDM based on pairwise Euclidean distances in the 2D frontoparallel plane, and a 3D geometric distance RDM based on Euclidean distanced in full calibrated 3D space. These models capture the representational structure expected if neural patterns reflect integrated spatial relationships rather than only individual feature dimensions. We tested these geometric distance representations in two complementary ways (Figure 5). First, we examined neural representational similarity to each of the 2D and 3D geometric distance models, while controlling for the other distance model, allowing us to identify neural sensitivity to each geometric structure. For comparison, the corresponding correlation-based RSA results, in which shared variance was not removed, are shown in the Supplementary (Figure S5). Second, because geometric-distance models can also share variance with individual spatial features, we repeated the analysis while additionally controlling for all ten individual component feature RDMs. This second analysis isolated unique geometric-distance representations that could not be explained by separable feature-level coding alone.

**Figure 5.**
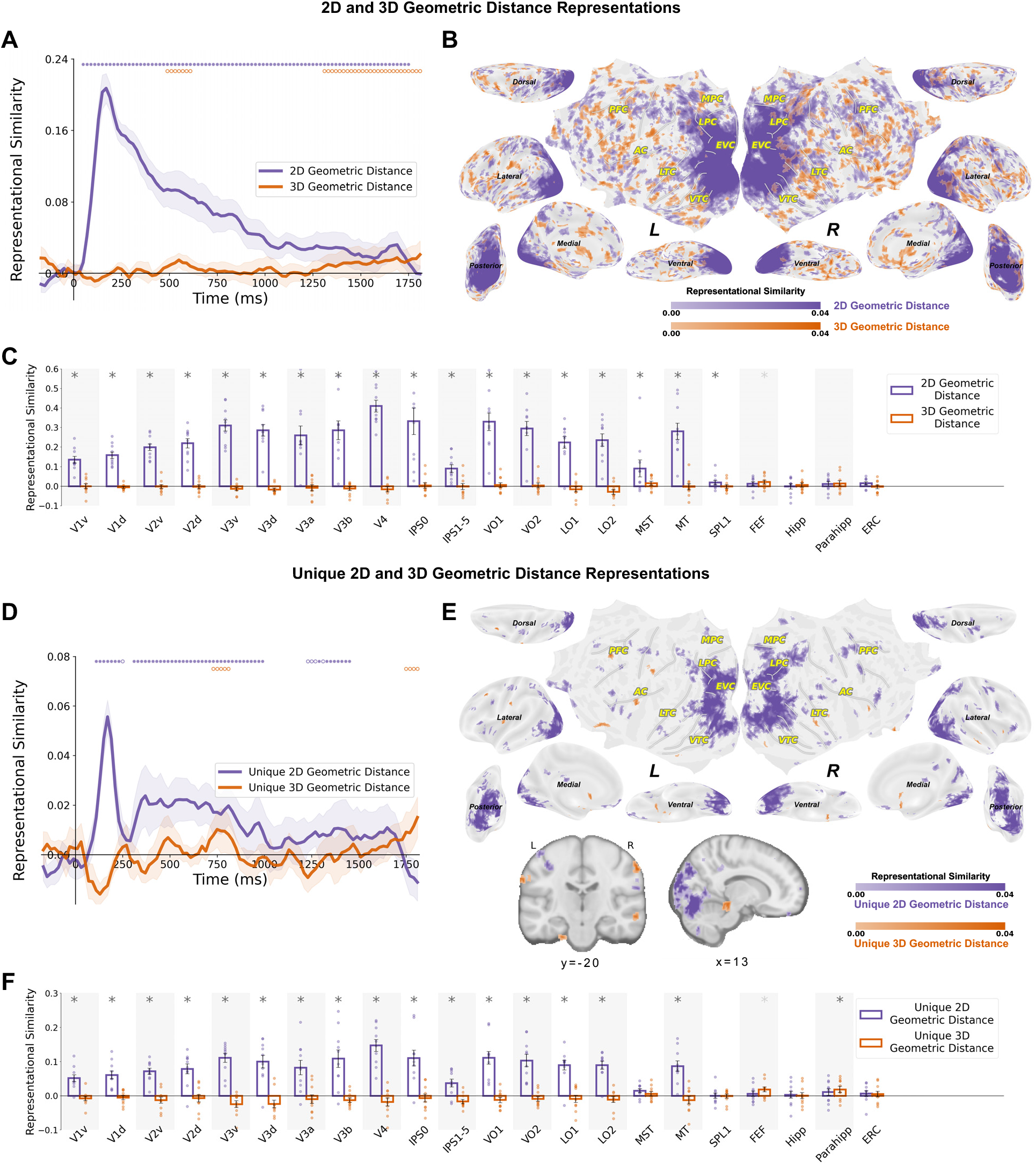
Spatiotemporal representations of 2D and 3D geometric distance. (A) Time-resolved representational similarity (partial Spearman correlation) between EEG temporal RDMs and the two geometric distance RDMs. Shaded areas indicate ±1 SEM across participants. Open circles indicate significant time points primarily driven by amplitude-based RDMs, whereas filled circles indicate significant time points primarily driven by pattern-based RDMs (permutation tests with correction for the number of hypothesis-based RDMs, followed by cluster-based correction, *p*<.05). (B) fMRI searchlight representational similarity (partial Spearman correlation) between fMRI temporal RDMs and the two geometric distance RDMs. fMRI searchlight representational similarity maps displayed on inflated cortical surfaces. Colored voxels indicate the searchlight centered on that voxel was significant (permutation tests with correction for the number of hypothesis-based RDMs, followed by cluster-based correction, *p*<.05). (C) ROI-based RSA results showing group-level representational similarity (partial Spearman correlation) of 2D and 3D geometric distances across 25 predefined ROIs. Light-gray asterisks indicate significant effects primarily driven by amplitude-based RDMs, whereas dark-gray asterisks indicate significant effects primarily driven by pattern-based RDMs (permutation tests with correction for the number of hypothesis-based RDMs, *p*<.05). Dots denote individual participants. Error bars indicate SEM. (D) Unique 2D and 3D geometric distance representations in EEG after controlling for all individual spatial feature RDMs (partial Spearman correlation). Shaded areas indicate ±1 SEM across participants. Open circles indicate significant time points primarily driven by amplitude-based RDMs, whereas filled circles indicate significant time points primarily driven by pattern-based RDMs (permutation test, cluster-corrected, *p*<.05). (E) Unique 2D and 3D geometric distance representations in fMRI after controlling for all individual spatial feature RDMs (partial Spearman correlation). fMRI searchlight representational similarity maps displayed on inflated cortical surfaces and anatomical slices. Colored voxels indicate the searchlight centered on that voxel was significant (permutation tests with correction for the number of hypothesis-based RDMs, followed by cluster-based correction, *p*<.05). (F) ROI-based RSA results showing group-level representational similarity (partial Spearman correlation) of unique 2D and 3D geometric distances across 25 predefined ROIs. Light-gray asterisks indicate significant effects primarily driven by amplitude-based RDMs, whereas dark-gray asterisks indicate significant effects primarily driven by pattern-based RDMs (permutation tests with correction for the number of hypothesis-based RDMs, *p*<.05). Dots denote individual participants. Error bars indicate SEM.

The first analysis is shown in Figure 5A-C. EEG analyses (Figure 5A) revealed robust encoding of both 2D and 3D geometric distances. Notably, 2D geometric distance representations emerged rapidly and remained significant throughout the epoch. In contrast, 3D geometric distance representations emerged later and were weaker than 2D geometric distance representations. fMRI analyses (Figure 5B-C) revealed a similar spatial dissociation: 2D geometric distance representations were broadly distributed across visual cortex, whereas 3D geometric distance representations were more spatially restricted and sparser, localized to patches in parietal, lateral occipital, and prefrontal regions. ROI-based analyses further confirmed this pattern, with robust 2D geometric distance effects across many visual ROIs and weaker 3D geometric distance effects in limited regions, such as FEF (Figure 5C). This spatiotemporal dissociation suggests a possible progression from planar to volumetric spatial integration in visual processing, analogous to the progression from 2D to 3D individual spatial feature dimensions.

We then asked whether these geometric-distance effects reflected integrated spatial structure beyond separable feature-level coding. To further isolate the unique contributions of these geometric distance representations beyond feature-level coding, we repeated the analysis above while additionally controlling for all ten individual spatial feature RDMs. This approach removes variance shared with feature-based coordinate representations, isolating the residual geometric integration component that may be otherwise masked by spatial features. EEG analyses (Figure 5D) revealed that unique 2D geometric distance representations persisted from early to late stages, whereas unique 3D geometric distance representations emerged only later. fMRI searchlight and ROI-based analyses (Figure 5E-F) revealed unique 2D geometric distance representations remained broadly distributed across visual cortex and adjacent regions. Nevertheless, unique 3D geometric distance representations were more spatially sparse and observed in limited regions, including patches near parahippocampal cortex – a region known to be involved in spatial navigation and scene representation (Aminoff et al., 2013; R. Epstein et al., 1999, 2003; R. A. Epstein, 2008).

Together, these findings demonstrate that neural coding of spatial location in human brain is not fully captured by independent feature dimensions. The human brain robustly represents integrated 2D geometric relationships which emerge early and are strongly and widely distributed, with additional, more restricted evidence for integrated 3D geometric structure which emerges later, weaker, and in more sparsely localized higher-order regions. For comparison, the corresponding correlation-based RSA results, in which shared variance was not removed, are shown in Supplementary Figure S6.

### Exploratory link between individual differences in depth perception and neural 3D representations

As an exploratory brain-behavior analysis, we next asked whether any of these spatiotemporal neural signatures of 3D visual processing might carry behavioral relevance in terms of 3D visual perception. Although our main EEG and fMRI task involved passive viewing of 3D stimuli, each participant completed the behavioral 3D cube adjustment task, allowing us to estimate a participant-specific depth magnitude gain from the mapping between binocular disparity and perceived depth. We therefore performed an exploratory analysis examining whether individual differences in depth perception were related to neural encoding of 3D-related information.

For each participant, we extracted the slope parameter *α* (depth magnitude gain) from the cube adjustment task, which reflects how strongly binocular disparity must be scaled to achieve a subjectively matched depth distance (perceived depth scaling). We then correlated this behavioral depth magnitude gain with each participant’s maximum neural representational similarity (across fMRI searchlight units; EEG had too few subjects to be meaningful for this analysis) for different neural measures including the depth- or 3D-related spatial features, 3D geometric distance and unique 3D geometric distance representations.

Among these measures, only the fMRI representation of the 3D radius feature (*r-3D*) showed a significant positive correlation with behavioral slope (*r*=.7917, *p*=.0064; Figure 6), indicating that participants with steeper disparity-to-depth mappings exhibited stronger neural encoding of 3D radius. All other correlations were non-significant (Figure S7), though we caution too much emphasis on this exploratory analysis of small sample size. That said, this finding suggests an intriguing potential link between individual differences in perceived depth magnitude and the strength of 3D spatial representations in the human brain.

**Figure 6.**
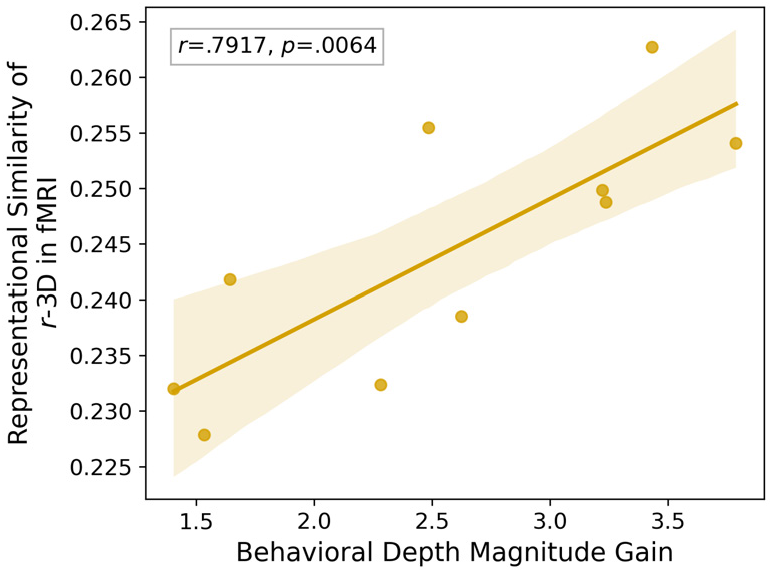
Correlation between behavioral depth magnitude gain and neural 3D radius encoding in fMRI. Each dot represents one participant. Shaded area indicates 95% confidence interval for the fitted regression line.

### Exploratory EEG–fMRI representational correspondence

As an additional exploratory analysis, we also examined the correspondence between EEG and fMRI representational structures to assess whether the spatiotemporal patterns observed in EEG align with spatial patterns in fMRI. Specifically, we computed representational similarity between time-resolved EEG RDMs and fMRI RDMs across cortical locations.

As shown in Figure 7, EEG-fMRI representational similarity revealed temporally evolving correspondence patterns. Earlier time stages showed stronger correspondence with posterior visual regions, whereas later time stages exhibited more distributed correspondence across higher-level cortical areas (Figure 7A). ROI-based analyses further quantified the dynamic similarity patterns (Figure 7B). Across regions, representational similarity generally peaked in the early post-stimulus period and gradually decreased over time. In addition, onset and peak latency analysis suggested a temporal progression across cortical regions, with earlier peaks in visual areas and later peaks in higher-order regions. These results provide converging evidence that the temporal dynamics observed in EEG are broadly consistent with the spatial organization of representations observed in fMRI.

**Figure 7.**
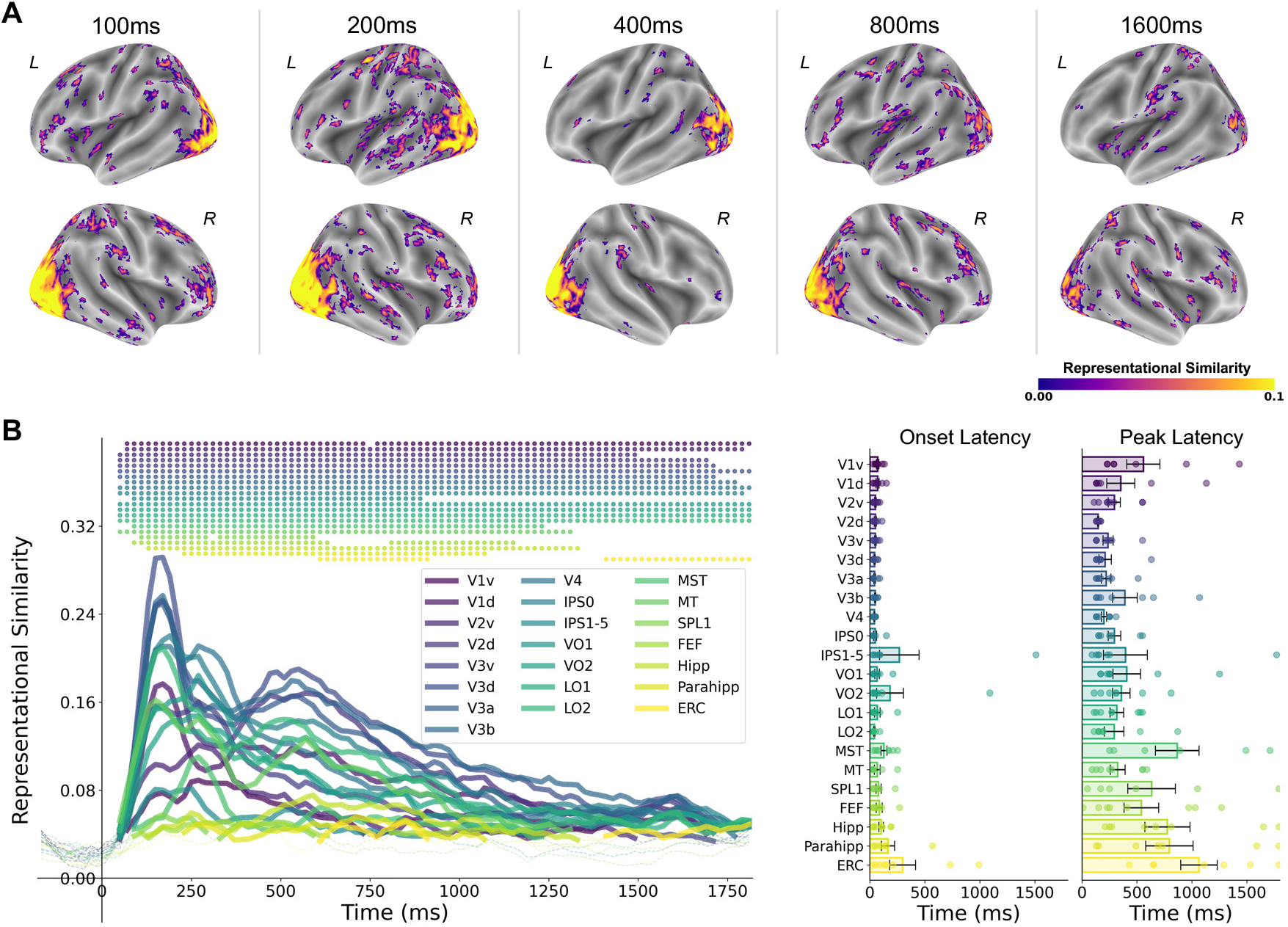
EEG-fMRI representational correspondence in 3D visual perception. (A) Cross-modal representational similarity between EEG RDMs at selected time points (100, 200, 400, 800, 1600 ms) and fMRI searchlight RDMs across cortex. Warmer colors indicate higher representational similarity. Colored voxels indicate the searchlight centered on that voxel was significant (permutation tests with cluster-based correction, *p*<.05). (B) Time-resolved EEG-fMRI representational similarity across ROIs. Left: Time courses of representational similarity for each ROI. Colored dots indicate significant timepoints (permutation tests with correction for the number of ROIs, followed by cluster-based correction, *p*<.05). Right: Onset and peak latency distributions across ROIs.

### Exploratory analyses of coordinate-format preferences across time and brain space

The feature-level analyses revealed that different spatial dimensions were not represented equally. In particular, 2D polar features such as *r* and *θ* showed stronger and earlier effects than 2D Cartesian features such as *x* and *y*, raising the possibility that neural spatial coding may be organized preferentially in some coordinate formats over others. We therefore performed exploratory analyses to ask whether neural representations showed evidence for specific coordinate formats across time and brain space, such as Cartesian coordinates defined by orthogonal axis-based dimensions or polar coordinates defined by radius and angular position (Figure 1B-D). We distinguished between “complete” coordinate representations, defined using a strict criterion that all individual constituent features within that coordinate system were significantly represented at that timepoint or cortical location, and “emergent” coordinate representations, defined using a more permissive criterion that captures coordinate-level preference when at least one coordinate-specific feature is represented (i.e., a feature uniquely associated with that coordinate system – e.g., among the 3D coordinate systems, *r* is specific to the Cylindrical system, while *x* and *y* are unique to the Cartesian system).

We first tested whether human neural activity reflects complete coordinate representations for any of these spatial coordinate systems (Figure 8A-B). In EEG data (Figure 8A), 2D polar coordinates showed robust and early representational similarity, emerging shortly after stimulus onset and remaining significant across extended time windows. In contrast, 2D Cartesian coordinates exhibited weaker and more temporally restricted effects. For 3D coordinate systems, complete representations were generally limited. Only 3D Cylindrical coordinates showed temporally restricted periods of significance, and this effect was weaker and less consistent compared to 2D representations. No 3D coordinate system showed sustained complete representations across time. The fMRI results showed a similar pattern (Figure 8B). Complete 2D coordinate representations of both 2D Cartesian and Polar were broadly distributed across visual cortex, but strikingly, the cortical topographies were largely dissociated. Polar representations were more spatially confined in early visual cortex and more foveal areas, with the Cartesian representations showing more widespread activation into peripheral visual areas and across higher-level dorsal and ventral visual pathway regions, extending into later temporal, parietal, and some prefrontal areas. In contrast, complete 3D coordinate representations were only significant for 3D Cartesian but spatially restricted. These findings suggest that under a strict “complete representation” criterion, neural activity within a given region or timepoint is more consistent with 2D coordinate frameworks than with fully specified 3D coordinate systems.

**Figure 8.**
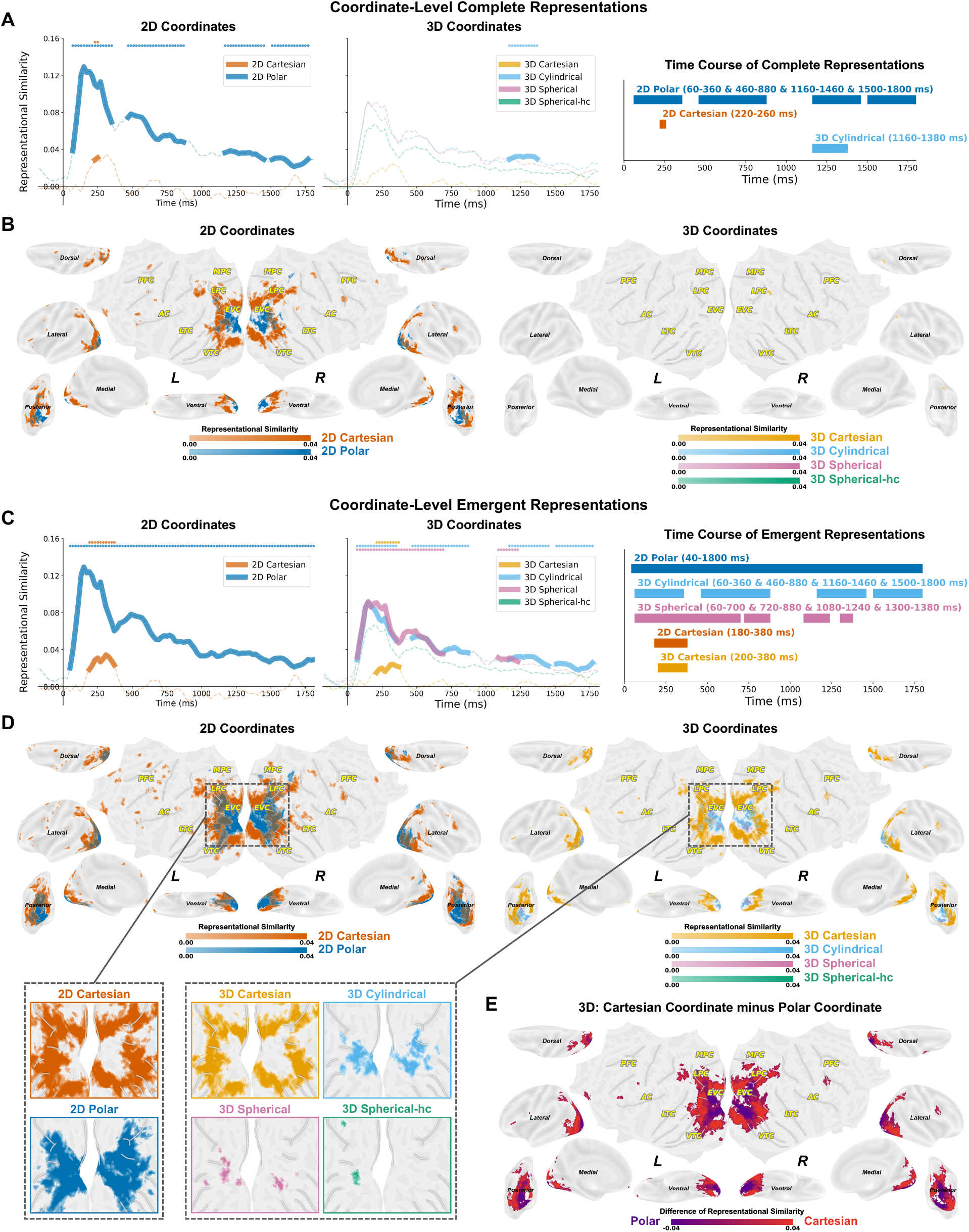
Complete and emergent coordinate-format representations. (A-B) Complete coordinate representations. Complete representations were defined using a strict criterion requiring all constituent spatial features within a coordinate system to show significant representational similarity (partial Spearman correlation). (A) EEG time courses and (B) fMRI searchlight maps of significant complete 2D and 3D coordinates. Thickened lines and Colored dots indicate significant timepoints (permutation test, cluster-corrected, *p*<.05). Voxels were labeled as significant if they showed representational similarity for all features in a certain coordinate system (permutation tests with correction for the number of hypothesis-based RDMs, followed by cluster-based correction, *p*<.05). (C-D) Emergent coordinate representations. Emergent representations were defined using a more permissive criterion capturing coordinate-level evidence when at least one coordinate-specific feature was significantly represented. (C) EEG time courses and (D) fMRI searchlight maps of significant emergent 2D and 3D coordinates. Thickened lines and Colored dots indicate significant timepoints (permutation test, cluster-corrected, *p*<.05). Voxels were labeled as significant if they showed representational similarity for all features in a certain coordinate system (permutation tests with correction for the number of hypothesis-based RDMs, followed by cluster-based correction, *p*<.05). (E) Direct comparison between Cartesian and Polar coordinate formats. (C) Differential maps showing Cartesian minus Polar representations for 3D coordinates (permutation tests with correction for the number of hypothesis-based RDMs, followed by cluster-based correction, *p*<.05). Here, we created a unified 3D Polar map by taking the voxelwise maximum of the emergent representations across 3D Cylindrical, 3D Spherical, and 3D Spherical-hc systems, and subtracted this from 3D Cartesian map.

We next examined emergent coordinate-level patterns using a more permissive criterion, asking whether neural activity reflects partial or feature-level evidence consistent with different coordinate formats (Figure 8C-D). In the EEG data (Figure 8C), this analysis revealed broader and more temporally extended coordinate-related effects. 2D polar representations remained dominant across much of the epoch, whereas 3D coordinate systems showed multiple temporally distinct periods of significant representational similarity. Especially, 3D Cylindrical and Spherical representations emerged earlier and more sustained over time than the 3D Cartesian representation. In the fMRI data (Figure 8D), emergent coordinate representations were more spatially distributed than complete representations. Different coordinate formats showed partially overlapping but heterogeneous spatial patterns across cortex, but these patterns were not entirely unstructured. To further characterize and summarize the organization of emergent 3D coordinate representations, we directly contracted 3D Cartesian and 3D Polar (including Cylindrical, Spherical, and Spherical-hc) coordinates (Figure 8E). This comparison suggested a broad spatial tendency, with posterior visual areas showing relatively stronger 3D Polar-coordinate representations and more anterior regions showing relatively stronger 3D Cartesian-coordinate representations. Thus, compared with the strict complete-representation analysis, the emergent analysis revealed richer and more spatially distributed coordinate-format representations, particularly for 3D coordinate systems, while also suggesting a coarse posterior-to-anterior organization in relative coordinate-format preferences.

## Discussion

In this study, we asked whether neural representations of 3D visual space are limited to separable component dimensions, such as 2D position and position-in-depth, or whether the human brain also encodes richer forms of 3D spatial structure. To address this question, we combined individualized perceptual calibration of binocular disparity, densely sampled 3D spatial stimulation, high temporal-resolution EEG, high spatial-resolution fMRI, and representational similarity analysis. This framework allowed us to place perceived depth on a common scale with horizontal and vertical position and to examine how 2D, depth/disparity-related, 3D feature-level, and geometric distance representations unfold across time and brain space.

Our results support two main conclusions. First, 3D spatial coding in the human brain showed a clear spatiotemporal hierarchy. 2D spatial representations emerged early and persisted strongly over time in EEG, and were broadly and robustly distributed across visual cortex in fMRI. Depth/disparity-related and 3D representations were later in time, weaker, and more spatially restricted, with sparser patches in primarily later brain areas. Second, and critically: neural coding of 3D visual perception extended beyond individual component feature dimensions. In other words, the neural mechanisms supporting 3D perception do not appear to operate simply by appending an independent depth signal onto existing 2D spatial maps. If that were the case, neural representational structure would be expected to be captured primarily by separable 2D position and signed position-in-depth dimensions. Instead, neural activity also contained information about certain 3D spatial features and integrated 3D geometric-distance structure. While these 3D representations were weaker and more restricted, the results overall suggest a transformation from dominant 2D coding toward more richly integrated spatial representations that may support perception of objects as embedded in 3D space. The exact nature of these 3D spatial representations remains to be explored, but the current study lays foundational groundwork and exciting theoretical advances for future work to build on.

Several methodological features of the present study were critical for making these analyses possible. First, we individually calibrated perceived depth using the cube-adjustment task. Many previous studies of disparity-defined depth reply on fixed physical disparity values across participants (Alvarez et al., 2021; Bridge & Parker, 2007; Finlayson et al., 2017). However, a fixed disparity does not necessarily correspond to the same perceived depth magnitude across individuals, making it difficult to compare depth directly with horizontal and vertical position. Our cube-adjustment task allowed us to estimate a participant-specific mapping between binocular disparity and perceived depth, enabling us to construct a calibrated 3D stimulus space for each participant. This calibration was essential for testing whether neural representations extend beyond 2D and depth component dimensions to higher-order integrated 3D spatial structures. Second, we densely sampled 64 locations across calibrated 3D space and combined high-temporal-resolution EEG with high-spatial-resolution fMRI, which enabled us to disentangle overlapping feature representations and robustly analyze their contributions in human brains. This comprehensive feature space allowed us to consider multiple types of 3D geometric relationships rather than only comparing 2D and depth axes, allowing for a more complete assessment of how the brain encodes spatial information. Finally, our use of partial correlation RSA framework enabled us to isolate the unique contributions of each spatial feature or geometric distance structure, going beyond standard correlation-based approaches that may conflate shared variance among features and misestimate the representational patterns (see correlation-based RSA results in Figure S4–S6). This was particularly important for interpreting depth-related and 3D effects: because the absolute disparity magnitude model (|*z*|) was included, significant signed-depth and 3D feature effects cannot be explained simply by differences in overall binocular disparity magnitude. Similarly, unique 3D geometric-distance effects were tested after removing variance shared with the component feature RDMs, including both depth and disparity magnitude.

These methodological advances allowed us to extend previous work on 2D and depth representations in visual cortex. Prior studies have characterized 2D spatial coding in terms of retinotopic position, eccentricity, polar angle, and related spatial dimensions (J. M. Carlson et al., 2011; Golomb & Kanwisher, 2012; Graumann et al., 2022; Kravitz et al., 2010), and other work has shown sensitivity to position-in-depth information conveyed by binocular disparity (Backus et al., 2001; Ban et al., 2012; Cumming & Parker, 1997; Henderson et al., 2019; Hibbard et al., 2017; Masson et al., 1997; Neri et al., 2004; Uka & DeAngelis, 2006; Welchman et al., 2005). A few recent studies have examined 2D position and binocular-disparity-defined position-in-depth representations in conjunction, but have done so using fixed disparity values or categorical depth levels, without estimating how those disparity values correspond to perceived depth magnitude for individual participants (Alvarez et al., 2021; Bridge & Parker, 2007; Finlayson et al., 2017). In contrast, our cube-adjustment task provided a participant-specific disparity-to-depth mapping, allowing perceived depth distances to be placed on the same perceptual metric scale as horizontal and vertical distances. Consistent with previous studies, we observed a hierarchical pattern of 2D to depth processing in the human brain. Indeed, we showed in a supplementary analysis a convincing replication of Finlayson et al., 2017’s analysis (correlation-based MVPA instead of partial correlation-based RSA) and results (spatial transition from 2D (the average of x and y) to depth (z) along the visual hierarchy; Figure S2). Crucially, our study extends these findings in several important ways. First, our use of EEG in addition to fMRI allowed us to uncover the detailed temporal processing trajectories of these different types of spatial information and link the temporal emergence of neural representations to patterns along the cortical visual hierarchy, producing a much richer view of neural processing. Second, we found that the human brain represents 3D visual space not only by encoding position-in-depth as an additional dimension alongside 2D location, but also by constructing richer forms of 3D spatial structure.

A key finding that neural coding beyond position-in-depth was detectable at the level of individual 3D features such as 3D radius and 3D polar angle, which previous studies have never examined. The results pattern suggests that different 3D feature representations are not strongly overlapping in time or brain space. Rather, they appear to be represented at different time points and in different cortical regions. This heterogeneity helps explain why averaged 3D effects were weak in the grouped analyses, while feature-level analyses nevertheless revealed significant 3D-related representational similarities.

The geometric-distance results further suggest that neural coding of 3D perception is not fully captured by separable feature dimensions. A representation based only on separable dimensions can encode where a stimulus falls along individual axes, but it does not necessarily encode the geometric relationships among locations. By constructing 2D and 3D geometric distance RDMs, we tested whether neural patterns reflected the pairwise spatial distances among locations. While 2D geometric distance was broadly represented, 3D geometric distance emerged only in specific regions, such as the parahippocampal cortex, and at middle and late timepoints in EEG. Interestingly, the parahippocampal cortex exhibited significant unique 3D geometric distance representation only after controlling for feature-level RDMs (Figure 6E-F), but not in the initial analysis (Figure 6B-C). This pattern likely reflects the removal of noisy variance shared with low-level spatial features, revealing a residual high-level 3D integration signal in this region. Such a dissociation is consistent with the parahippocampal cortex’s proposed role in encoding abstract spatial relationships, such as real-world navigation and scene perception (Aminoff et al., 2013; R. Epstein et al., 1999, 2003; R. A. Epstein, 2008), beyond simple feature-based metrics.

Importantly, our geometric distance RDMs were designed to capture the spatial relationship among stimuli in terms of their relative locations – how far each stimulus is from the others – rather than their absolute positions with respect to the reference point. This finding aligns with previous studies emphasizing the brain is sensitive not only to absolute location but also to relative spatial information (Hayworth et al., 2011; Roth, 2016), which underlies functions such as spatial comparison, grouping, and navigation (Berens et al., 2021; Byrne et al., 2007; Roth, 2016; Uchimura et al., 2015). Together, our results may suggest that human brains encode not only feature-based spatial information but also higher-order spatial structure that could capture relational geometry among multiple objects in space.

A key consideration in interpreting the present results is the role of RSA in linking neural activity to representational hypotheses. RSA does not identify isolated parameters directly, but instead evaluates whether the similarity structure of neural responses is consistent with model-defined representational spaces. In the present study, this framework allowed us to test whether spatial coding is better characterized by separable component dimensions, such as 2D position and depth, or by more integrated geometric relationships among locations. In addition, our RSA analyses concern the distinction between amplitude-based and pattern-based RDMs. These two measures capture complementary aspects of neural coding. Amplitude-based RDMs reflect differences in overall response magnitude across conditions and may therefore be sensitive to univariate spatial tuning, response strength, or stimulus-driven salience. Pattern-based RDMs reflect differences in multivariate activity patterns across channels or voxels and may better capture distributed representational structure that is not reducible to mean response differences. We therefore chose not to interpret one RDM type as universally superior; instead, we treated them as complementary windows into different representational formats. Indeed, in the present study, some effects were primarily driven by amplitude-based RDMs whereas others were driven by pattern-based RDMs, indicating that 3D spatial coding is expressed through both response magnitude and distributed pattern structure.

Several exploratory analyses further contextualized these main findings. First, the brain-behavior analysis suggested a possible relationship between individual differences in perceived depth scaling and neural 3D spatial coding, with stronger behavioral depth magnitude gain associated with stronger neural encoding of 3D radius; however, this result should be interpreted cautiously given the small sample size. Second, the EEG-fMRI representational correspondence analysis linked the temporal dynamics observed in EEG with the spatial organization observed in fMRI, showing that earlier EEG patterns corresponded more strongly to posterior visual regions whereas later patterns showed broader correspondence with higher-order cortical regions. Third, the coordinate-level analyses provided an exploratory first step toward characterizing the coordinate formats in which these spatial codes may be expressed.

Building on these findings, an important open question concerns the reference frames that underlie these spatial codes. Our analyses primarily defined locations relative to a gaze-centered origin, consistent with retinotopic or egocentric representations, with the exception of the head-centered Spherical (Spherical-hc) coordinates. However, because participants maintained central gaze fixation, our paradigm does not directly dissociate retinotopic from spatiotopic encoding, nor egocentric from allocentric coding (Burgess et al., 2007; Byrne et al., 2007; Crespi et al., 2011; Duhamel et al., 1997; Filimon, 2015; Gardner et al., 2008; Golomb & Kanwisher, 2012; Klatzky, 1998; Zaehle et al., 2007). Nonetheless, the involvement of regions such as the parahippocampal cortex – known for allocentric and scene-based representations (Aguirre et al., 1996; Ekstrom et al., 2014; Parslow et al., 2004; Rolls, 2020; Zaehle et al., 2007) – raises the possibility that multiple reference frames may coexist across cortical systems. Future work could explicitly manipulate fixation or use immersive environments to clarify how these reference frames interact in supporting 3D spatial perception.

Another important open question is whether the spatiotemporal dynamics observed here generalize to other depth cues, such as motion parallax, texture gradients, relative size, shading, occlusion, and perspective, or to naturalistic settings in which multiple depth cues are integrated. Relatedly, our passive viewing paradigm was designed to characterize baseline 3D spatial representations under minimal task demands, but 3D perception in natural behavior likely involves attention, action planning, eye movements, memory, and predictive processing. Future studies combining calibrated 3D stimulation with active viewing, task manipulations, immersive environments, or memory-guided navigation could test how these processes shape the emergence and use of 3D spatial representations.

Beyond the specific findings reported here, we believe that a broader impact of this project lies in the resources and methodological framework it offers. To our knowledge, this study introduces the first large-scale, multimodal neuroimaging dataset for 3D spatial perception, and we provide a first and novel step in investigating the spatiotemporal neural representations of 3D visual perception in the human brain. Although recent advances in large-scale fMRI and EEG datasets have opened new possibilities for investigating the neural basis of visual cognition (Allen et al., 2022; Chang et al., 2019; Gifford et al., 2022; Grootswagers et al., 2022; Hebart et al., 2023), no such dataset currently exists for 3D spatial perception. We have publicly released our data, which we hope will serve as a foundation for future research efforts – not only to further advance our understanding of the neural mechanisms underlying 3D perception, but also to support translational applications such as decoding spatial location from brain activity in brain-computer interface (BCI) contexts across both cognitive neuroscience and bioengineering domains. Importantly, by making this dataset openly available, we also aim to enable the community to test alternative hypotheses and different models of 3D space representation, including approaches or feature dimensions beyond those we have explored here. In addition to the dataset, our work offers a generalizable methodological framework that integrates individual perceptual calibration, dense 3D spatial sampling, and multimodal RSA to probe high-resolution spatial encoding across both time and brain space.

Several methodological considerations should be acknowledged. First, the number of participants was modest, which may affect group-level sensitivity, particularly for time-resolved EEG analyses and feature-level comparisons. However, this limitation should be interpreted in light of the study’s dense within-subject sampling in the present design, which provides repeated measurements across a high-dimensional stimulus space and can improve the reliability of condition-level representational estimates (Allen et al., 2022; Gratton & Braga, 2026; Hebart et al., 2023; Naselaris et al., 2021). Second, although eye position was monitored online to ensure fixation, the eye-tracking data were not retained for offline trial-wise analysis, and the monocular eye-tracking setup did not permit direct quantification of vergence eye movements. Although fixation was actively controlled during acquisition – particularly in EEG, where fixation breaks triggered immediate trial abortion—and the nearest stimulus locations remained relatively peripheral (~2.12° eccentricity), we cannot fully rule out residual contributions from very small eye movements such as micro-saccades or from vergence-related processes (Quax et al., 2019). Importantly, participants performed a fixation-change detection task at central fixation, with the fixation target presented at zero disparity, encouraging them to maintain fixation and vergence near the central depth plane rather than tracking the depth of the peripheral stimuli. Thus, while eye-movement and vergence-related factors remain a potential limitation, they are unlikely to account for the spatiotemporal pattern of dissociable 2D, depth, and 3D-related representations observed here.

The exploratory coordinate-format results should also be interpreted in light of our stimulus sampling scheme. The 64 stimulus locations were uniformly sampled in Cartesian coordinates, which enabled systematic coverage of calibrated 3D space but did not account for eccentricity-dependent cortical magnification, whereby foveal and peripheral locations are represented nonuniformly across visual cortex (Deyoe et al., 1996; Dougherty et al., 2003; Harvey & Dumoulin, 2011; Sereno et al., 1995; Wandell et al., 2007). This is particularly relevant for interpreting differences between Cartesian and polar representations, because polar features such as radius/eccentricity and polar angle may be influenced by the distribution of sampled locations across the visual field and by the nonuniform cortical representation of foveal versus peripheral space. To mitigate this issue, our RSA computations were based on Spearman partial correlations, a rank-based method that is robust to differences in scale and nonlinearity, and controls for shared variance across all feature models. This approach could minimize the influence of global activation magnitude and cortical surface area differences. Nevertheless, the cortical magnification may still lead to underestimation or distortion of representational similarity results in certain regions. Future work using eccentricity-informed or alternative spatial sampling schemes could therefore help clarify whether the apparent differences between Cartesian and polar coordinate-format representations reflect intrinsic neural coding preferences, properties of the sampled stimulus space, or both.

Finally, an important direction for future research concerns how task demands modulate spatial encoding. In the current study, we employed a passive viewing paradigm to establish a baseline map of 3D spatial representations under minimal cognitive engagement. However, 3D perception likely involves feedback and predictive processes, including expectations about spatial structure. Because the present paradigm does not dissociate feedforward and feedback contributions, future studies combining task manipulations, temporal modeling, or model-based approaches will be needed to determine how predictive processes contribute to the emergence of depth-related and 3D spatial representations. Future work can build on our methodological framework to investigate coordinate-system dynamics in active viewing, naturalistic environments, or memory-guided navigation. These extensions will help us clarify not only why human brains apply different systems to represent spatial locations, but also how 3D spatial representations interface with eye movements, motor control, attention, and decision-making processes.

## Methods

### Participants

Ten participants participated in the study (7 females and 3 males, mean age = 26.0 ± 5.5 years) for monetary compensation ($15/hr for the behavioral session, and $20/hr for the fMRI and EEG sessions). Eight participants each completed one behavioral session, two EEG sessions, and two fMRI sessions in five separate days. The other two participants completed one behavioral session and two fMRI sessions in three separate days. All participants first completed the behavioral session, and subsequently did the neuroimaging sessions, scheduled according to participant availability and the scanner and EEG lab calendars. All participants had normal or corrected-normal vision, and were prescreened for MRI eligibility. The study protocol was approved by The Ohio State University Biomedical Sciences Institutional Review Board.

Our sample size was determined based on prior neuroimaging studies using large-scale EEG and/or fMRI to investigate high-dimensional neural encoding (Allen et al., 2022; Chang et al., 2019; Gifford et al., 2022; Hebart et al., 2023). Given the within-subject design and the large number of trials per participant across five sessions, the dataset affords high sensitivity to spatiotemporal representational structure at both individual and group levels. While a larger sample may improve generalizability (Allen et al., 2022; Gratton & Braga, 2026; Hebart et al., 2023; Naselaris et al., 2021), our findings reflect robust effects within and across participants and provide a strong foundation for future replication and extension studies.

### General Experimental Setup

Dynamic random dot stereogram stimuli (RDS) were generated using Psychtoolbox extension (Brainard, 1997) for MATLAB (Math Works). Depth from binocular disparity was achieved using red/green anaglyph glasses paired with Psychtoolbox’s stereomode. For all the experiments in our study, we created a 3D space composed of a low-contrast, large background field (12° × 12°) placed at the central depth plane of the screen, framed with additional depth cues making a 3D reference frame. The frontmost and backmost depth frames were rendered with disparities of +20 arcmin and −20 arcmin, respectively, creating a 40 arcmin total depth range. The background field consisted of static random dot stimuli (RDS; 5 dots/deg^2^) comprised of light and dark gray dots (each sized 0.18° × 0.18°) on a mid-gray background. To enhance the 3D percept and provide a stable spatial reference across depth, we also displayed visual depth cues composed of vertical and horizontal grid lines framing the 3D space outside the stimulus area. This grid formed a perspective-like scaffold, giving observers a consistent visual structure suggestive of depth to encourage perception of a 3D space. The main experimental stimuli were cubes of high-contrast dynamic RDS (each cube sized 2.4° × 2.4° × 2.4°, composed of 0.18° × 0.18° white and back grey dots, with 10 dots/deg^2^), presented at different locations within this 3D space, as described in the sections below.

For behavioral and EEG sessions, stimuli were presented on a 21-inch LCD monitor (resolution 1920 × 1080 at 240 Hz), and participants were seated at a chinrest 74 cm from the monitor. For fMRI sessions, stimuli displayed with a DLP projector onto a screen mounted in the rear of the scanner (resolution 1280×1024 at 60 Hz), and participants viewed from a distance of 74 cm via a mirror above their heads attached to the head coil.

### Behavioral Session

The behavioral session included three tasks. The first two tasks were used to confirm that the participants could accurately perceive and discriminate different depth distances with RDS from binocular disparity, and the third task was used to measure different disparity parameters corresponding to different depth distances. In addition, the extensive exposure during the whole behavioral session served to acclimate participants to the depth information in these displays, enhancing their sensitivity to depth cues in the subsequent main tasks.

In Task 1 – Single stimulus depth judgement task, participants viewed a single 2.4° × 2.4° square patch of dynamic RDS presented at the 2D center of the screen, appear either in front or behind the screen (fixation) plane. The stimulus depth was set to one of four disparity levels: +15, +5, −5, −15 arcmin (relative to the fixation depth plane). Participants were asked to judge whether the stimulus appeared in front of (closer to them) or behind (further away) the screen. Each trial was self-paced with no time limit and no eye movement restrictions. After participants responded, feedback indicating “correct” or “incorrect” was displayed, and the trial ended. Participant completed four blocks in total, each consisting of 24 trials (4 depth levels × 6 repetitions), with trials presented in a fully randomized order. After completing the first two blocks, participants reversed the orientation of their glasses (i.e., left-red/right-green or left-green/right-red). In one block per orientation, the key mapping was “A” for “in front” and “L” for “behind”, while in the other, the mapping was reversed (“L” for “in front”, “A” for “behind”).

In Task 2 – Two-stimulus depth judgment, participants viewed two dynamic RDS patches presented simultaneously to the left and right of central fixation (each 2.4° × 2.4°, with center positions 3° horizontally from fixation). The two RDS patches were presented at different depth levels, forming one of six predefined disparity pairs: left: −15 arcmin / right: −5 arcmin; left: −5 arcmin / right: −15 arcmin; left: −5 arcmin / right: +5 arcmin; left: +5 arcmin / right: −5 arcmin; left: +5 arcmin / right: +15 arcmin; left: +15 arcmin / right: +5 arcmin. The two stimuli could both appear in front of, behind, or straddling the fixation (screen) depth plane. Participants were instructed to judge the relative depth of the two stimuli. At the beginning of each block, participants were informed whether they should report which stimulus was closer or which was farther. For example, in a “closer” block, participants pressed the ‘A’ key if they thought the left stimulus was closer, or the ‘L’ key if they thought the right stimulus was closer. Each trial was self-paced with no time limit and no restriction on eye movements. Once the participant responded, feedback (“correct” or “incorrect”) was displayed, and the trial ended. Participants completed four blocks, each consisting of 12 trials (6 depth pairs × 2 repetitions), presented in randomized order. After two blocks, the color orientation of the anaglyph glasses was reversed (left-red/right-green or left-green/right-red; counterbalanced across participants). For each anaglyph orientation, one block required judging which stimulus was closer, and the other required judging which was farther.

All our ten participants showed > 90% accuracy in both Task 1 and 2, which verified that they were sensitive to the depth triggered by binocular disparity. Then they did the Task 3.

In Task 3 – cube adjustment task, participants viewed four RDS patches (each 2.4° × 2.4°) presented simultaneously in the upper-left, upper-right, lower-left, and lower-right quadrants of the screen. These four stimuli corresponded to the four vertices of a square shape defined by their farthest edges from fixation. Across trials, the horizontal and vertical distances of this “virtual” square were simultaneously manipulated to form one of 11 physical sizes: 5.4°×5.4°, 6.0°×6.0°, 6.6°×6.6°, 7.2°×7.2°, 7.8°×7.8°, 8.4°×8.4°, 9.0°×9.0°, 9.6°×9.6°, 10.2°×10.2°, 10.8°×10.8°, or 11.4°×11.4°. Crucially, the four stimuli were arranged such that two diagonally opposite patches were presented in front of the fixation plane and the other two behind the fixation plane, with equal absolute disparities (i.e., symmetrical distance from fixation). The initial disparity was randomly selected on each trial. Participants performed a cube adjustment task: using the up/down arrow keys, they simultaneously moved the front and back planes either closer to or farther from the fixation plane, until they perceived the four vertices as forming a cube in 3D space. Note that while depth positions varied, the 2D square size on the screen remained fixed within each trial. Each trial was self-paced with no time limit and no restriction on eye movements. Participants pressed the “Enter” key to confirm when they believed the configuration matched a cube, thus ending the trial. Participants completed two blocks of 66 trials each (11 horizontal/vertical square sizes × 2 depth arrangements (half with top-left and bottom-right stimuli in front, and half with them behind) × 3 repetitions), with trials presented in random order. After the first block, the orientation of the anaglyph glasses was reversed (left-red/right-green or left-green/right-red). This task allowed us to determine the individualized perceptually-matched binocular disparities corresponding to a range of reference distances from the fixation plane, ranging from 2.7° to 5.7° in 0.3° increments (i.e., half of the cube’s front-back extent). These distances reflect the amount of disparity required for a specific participant to perceive the four stimuli as forming a cube in 3D space, providing a participant-specific calibration of depth perception in stereoscopic space, allowing us to match the depth and 2D distances sampled in the main tasks.

### Individualized Depth Distance Quantification

To establish a participant-specific mapping between perceived depth distance and binocular disparity, we analyzed data from the cube adjustment Task (Task 3). We fit a linear function to each participant’s data using least-squares regression, with perceptual matched disparity as the dependent variable and reference distance as the independent variable. Given that a depth distance of 0° should yield zero disparity, we constrained the fit to pass through the origin, yielding a function of the form *Y*_disparity_ = *αX*_reference_distance_. The slope *α* captures the participant-specific transformation from reference distance (in degrees of visual angle) to perceptually-matched binocular disparity (in arcmin). This slope can be interpreted as an individualized “depth magnitude gain”, indexing how strongly binocular disparity must be scaled to achieve a subjectively matched depth distance (perceptual depth scaling). Figure S1 shows the model fitting results for all ten subjects.

This individualized linear model was used to compute the disparity corresponding to any desired depth distance for each participant, enabling precise, perceptually calibrated depth manipulation across all subsequent experimental sessions.

### EEG Session

In each EEG session, during each trial, participants viewed a small dynamic RDS cube stimulus (2.4° × 2.4° × 2.4°) presented at one of 64 possible 3D spatial locations. The 64 locations were defined by a 4 × 4 × 4 (horizontal × vertical × depth) grid with fixation at the origin ([0° 0° 0°]). The 64 possible stimulus locations were the intersections at the following grid coordinates: horizontal *x* position: −4.5°, −1.5°, 1.5°, 4.5° × vertical *y* position: −4.5°, −1.5°, 1.5°, 4.5° × depth *z* position: −4.5°, −1.5°, 1.5°, 4.5°. The exact amount of binocular disparity required to produce each depth position was individually calibrated for each participant using the linear function derived from the cube adjustment task (behavioral Task 3). The stimulus was a cube centered on these locations, and our design of the cube size and spacing was to ensure that stimuli at adjacent locations were spatially independent, with no overlap in 3D space. Such separation was critical for disentangling neural activity patterns associated with individual spatial locations.

Each stimulus was presented for 1.5 s, followed by a variable interstimulus interval (ISI) of 0.5-0.7 s (uniformly jittered). Participants were instructed to maintain central fixation at all times. The fixation cue consisted of two concentric dark gray circles (outer diameter: 0.45°; inner diameter: 0.15°) with a light gray crosshair at the center, presented at coordinates [0,0,0]. To ensure attention and maintain fixation, participants performed a fixation-change detection task: on a small proportion of trials, the central cross briefly changed into an “X” shape, and participants were instructed to press the spacebar upon detecting the change. Participants completed 18 blocks in each session, each consisting of 128 trials (64 spatial locations × 2 repetitions), with trials presented in fully randomized order. To control for any low-level color or contrast differences between eyes or monocular-based cues, participants reversed the orientation of the red/green anaglyph glasses across sessions, such that the red filter was on the left eye for once session and the right eye for the other. The order of initial color assignment was counterbalanced across participants. This flip manipulation counterbalanced which eye received the red versus green images across sessions, eliminating simple monocular or color-based cues. Importantly, near and far positions had equal disparity magnitudes.

### fMRI Session

In each fMRI session, participants completed ten to twelve main task runs (6.1 mins per run). Similar to the EEG blocks, each run consisted of 128 trials (64 spatial locations × 2 repetitions). On each trial, a dynamic RDS cube stimulus (2.4° × 2.4° × 2.4°) was presented for 1.5 s at one of the 64 spatial locations (same exact stimulus positions and presentation time as EEG task). Interleaved with the 128 stimulus trials were 42 blank trials (no stimulus presented), inserted randomly such that no more than two consecutive blank trials occurred, to achieve sufficient jitter for modeling rapid event-related fMRI BOLD activity. Participants were instructed to maintain fixation at the center of the screen and perform a fixation change detection task, pressing the spacebar whenever the central fixation symbol changed from a “+” to an “x”; these fixation changes could occur on blank trials as well as stimulus trials.

In addition to the main task runs, participants also completed two retinotopic mapping runs using standard rotating wedge stimuli (Engel et al., 1994; Sereno et al., 1995). High-contrast radial checkerboard patterns were presented as 60° wedges and flickered at 4 Hz. Maximal eccentricity was 16° and the central 1.6° foveal region was not stimulated (except for a central fixation point). One run rotated clockwise, and the another run rotated counter-clockwise through 7 cycles with a period of 24 s/cycle. During these runs, participants fixated at the center of the display and presses a button every time the black fixation dot dimmed to gray.

As with the EEG sessions, the anaglyph glasses direction was reversed across fMRI sessions, with the order randomized across participants.

### Eye Tracking

Monocular eye position was monitored with an EyeLink 1000 eye-tracking system in both EEG lab and the fMRI scanner. The eye tracker was calibrated using a nine-point grid method at the beginning of each neuroimaging session and re-calibrated as necessary.

In the EEG sessions, eye position was continuously monitored online. Whenever gaze deviated by more than 1.5° from central fixation during the 1.5 s stimulus period, the trial was immediately terminated, an on-screen message reminded the participant to maintain fixation (“please fixate at the center”), and that trial was subsequently reinserted into the experiment in randomized order. Thus, all EEG data come from trials in which central gaze position was verified.

In the fMRI sessions, fixation was also monitored online throughout each run, but trials were not terminated or removed for gaze deviations to preserve the fMRI trial timing and design. Because quantitative calibration and stable pupil tracking were occasionally less reliable in the scanner environment during stereoscopic viewing through anaglyph glasses, fixation quality was assessed primarily through real-time monitoring of the eye-tracking display by the experimenter, as well as through accurate performance on the fixation detection task. Runs showing obvious fixation instability during stimulus presentation were excluded from further analysis. However, the eye-tracking signals were not stored in a form suitable for more sensitive retrospective trial-wise exclusion analyses.

### EEG Acquisition

EEG experiments were carried out at the EEG lab in Department of Psychology at The Ohio State University. EEG data were recorded using an elastic cap (Brain Products ActiCap) with 64 active electrodes (including one online reference channel: FCz, and other 63 channels: FP1, FP2, AFz, AF3, AF4, AF7, AF8, Fz, F1, F2, F3, F4, F5, F6, F7, F8, FC1, FC2, FC3, FC4, FC5, FC6, FT7, FT8, FT9, FT10, Cz, C1, C2, C3, C4, C5, C6, T7, T8, CPz, CP1, CP2, CP3, CP4, CP5, CP6, TP7, TP8, TP9, TP10, Pz, P1, P2, P3, P4, P5, P6, P7, P8, POz, PO3, PO4, PO7, PO8, Oz, O1, O2) arranged in the standard 10-20 layout, and a BrainVision actiCHamp amplifier at a sampling rate of 1000 Hz with the online filtering (between 0.1 Hz and 100 Hz). Electrode impedances were reduced to <20 kΩ before the commencement of each experiment session.

### fMRI Acquisition

fMRI experiments were carried out at The Ohio State University Center for Cognitive and Behavioral Brain Imaging with a Siemens Prisma 3T MRI scanner using a 32-channel phase array receiver head oil. Functional data were acquired using a T2-weighted gradient-echo planar imaging (EPI) sequence (TR = 2000 ms, TE = 30 ms, flip angle = 72°; 2 × 2 × 2 mm voxel size; 72 axial slices; no gap). The acquisition was aligned to the anterior commissure-posterior commissure (AC-PC) plane. Multiband acceleration was applied using the CMRR mbep2d_bold sequence with a multiband factor of 3. A high-resolution T2-weighted turbo spin echo (TSE) sequence was acquired for hippocampal subfield segmentation (TR = 4800 ms, TE = 106 ms, flip angle = 135°; voxel size = 0.5 × 0.5 × 2 mm). Also, we collected a high-resolution MPRAGE anatomical scan (1 mm^3^) for each participant.

### EEG Preprocessing

Offline EEG preprocessing was conducted in Python, using the MNE-Python package (Gramfort et al., 2013) along with customized scripts based on a previously published pipeline (Lu et al., 2024). EEG data from two recording sessions for each participant were merged into a single dataset. A band-pass filter from 0.03 to 50 Hz was applied, and independent component analysis (ICA) was used to identify and remove artifacts related to eye blinks and eye movements (Drisdelle et al., 2017; Jung et al., 2000). Signals from channels TP9 and TP10, placed over the left and right mastoids, were used for re-referencing. The continuous EEG data were resampled to 50 Hz using standard anti-alias filtering then segmented into epochs from −200 ms to 1800 ms relative to stimulus onset. Baseline correction was performed by substracting the mean voltage of a 100 ms pre-stimulus period (−100 to 0 ms) for each trial and channel. This resulted in a matrix of preprocessed EEG data for each participant with dimensions 4,608 trials × 63 channels × 100 timepoints.

To obtain location-specific event-related potentials (ERPs), we averaged the 72 repeated trials corresponding to each of the 64 stimulus location labels. This procedure yielded an ERP matrix with dimensions 64 stimulus locations × 63 electrode channels × 100 timepoints for each participant, which was used for all subsequent analyses.

### fMRI Preprocessing

fMRI data were preprocessed using Brain Vogager QX (Brain Innovation). Preprocessing included slice timing correction, head motion correction, temporal filtering, and normalization to Talairach space (Talairach & Tournoux, 1988). No spatial smoothing was applied to the data used for representational similarity analysis. A whole-brain random-effects general linear model (GLM) was run for all main task runs across fMRI sessions to calculate beta weights for each voxel, for each spatial location condition, for each participant. This yielded a beta-weight matrix for each participant with dimensions 64 stimulus locations × 432,216 voxels. Each participant’s cortical surface for each hemisphere was inflated and flattened into cortical surface space for retinotopic mapping.

Retinotopic regions of interest (ROIs) including V1v, V1d, V2v, V2d, V3v, V3d, and V4 were functionally defined based on individual retinotopic mapping data. Additional ROIs—V3a, V3b, IPS0, IPS1–5, VO1, VO2, LO1, LO2, MST, MT, SPL1, and FEF—were defined using maximum probability maps from the Wang et al. probabilistic atlas (Wang et al., 2015). Medial temporal lobe ROIs, including the parahippocampal cortex, hippocampus, and entorhinal cortex (ERC), were anatomically defined using individual subject segmentations provided by FreeSurfer (Fischl, 2012).

### Representational Similarity Analysis

Our key analyses were based on representational similarity analysis (Kriegeskorte et al., 2008). The subsections below explain the details of how we computed representational dissimilarity matrices (RDMs) based on hypothetical representational spaces and actual neural signals (fMRI or EEG), and how we measured the unique representation of each spatial feature and geometric structure using the partial correlation approach (Dobs et al., 2019; Lu & Golomb, 2024). All analyses were implemented using customized code adapted from the NeuroRA toolbox (Lu & Ku, 2020).

#### Hypothesis-based Representational Dissimilarity Matrices (RDMs)

We constructed ten hypothesis-based spatial feature RDMs, each reflecting a distinct spatial feature dimension (*x, y, z, |z|, r, θ, r-3D, Φ, r-3D-hc, Φ-hc)*, plus two additional hypothesis-based geometric distance RDMs reflecting the combined geometric relationships among different spatial locations in 2D or 3D space (Figure 1E).

For the *x*, *y*, and *z* RDMs, we extracted the respective Cartesian coordinates (horizontal, vertical, and depth positions) of the 64 stimulus locations along the X-, Y-, and Z-axes. Each RDM was constructed as a 64 × 64 matrix where each cell reflects the dissimilarity along the given spatial dimension between a pair of two stimulus positions. For example, to compute the spatial dissimilarity between a stimulus centered at position #18 (*x*=−1.5°, *y*=−4.5°, *z*=−1.5°) and a stimulus centered at position #51 (*x*=4.5°, *y*=−4.5°, *z*=1.5°), we would calculate the absolute differences in their x, y, and z locations, respectively. This would produce a dissimilarity value of 6 for that cell of the X-RDM, a 0 for the Y-RDM, and a 3 for the Z-RDM. These hypothesis-based RDMs reflect the representational similarity patterns expected if the brain represented purely horizontal (*x*), vertical (*y*), or depth (*z*) information.

The z dimension reflects spatial position along the signed z-axis, ranging from stimuli perceived as appearing increasingly behind the fixation plane (negative) to stimuli appearing increasingly in front of the fixation plane (positive). Because depth here is cued by binocular disparity, it is important to distinguish perceptual (signed) depth-related coding from effects driven by absolute disparity magnitude, so we included a *binocular disparity magnitude* (|*z*|) RDM as a control. For this RDM, we extracted the absolute value of depth distance |z| of the 64 stimulus locations (corresponding to the magnitude of binocular disparity between the two eyes) and then computed the differences in |z| magnitude between each pair of stimulus locations; e.g., the example stimulus pair above would produce a 0 for |z|-RDM.

For the *r* and *θ* RDMs, we analogously extracted the respective Polar coordinates (radius and polar angle) of the 64 stimulus locations along the r- and *θ*-axes. For *r,* we computed the 2D radial distance from each location to the fixation point on the screen. For *θ*, we calculated the corresponding polar angle relative to the positive X-axis within the 2D plane. For each of these, the RDMs were computed as the absolute difference in radius or polar angle between each pair of stimulus conditions.

Analogous to how 2D space can be represented in Cartesian or Polar coordinates, 3D space can be represented in 3D Cartesian (*x, y, z*), 3D Polar Cylindrical (*r, θ, z*), or 3D Spherical (*r-3D, θ, Φ)* coordinates (Figure 1). For the *r-3D* feature, we computed the 3D radial distance from each stimulus location to the fixation point centered on the screen in the 3D space. For the *Φ* feature, we calculated the angle between this 3D direction vector and the positive Z-axis. We also calculated versions of these features assuming a spherical representation centered on the head instead of the screen. For the *r-3D-head-centered* feature and the *Φ-head-centered* feature, we computed the 3D distance and angle from each location to the participant’s head position as the reference point in the 3D space. For each of these features, dissimilarity values in the RDM were defined as the absolute differences in these dimensions between each pair of stimulus conditions.

Additionally, we constructed two additional hypothesis-based geometric distance RDMs, each reflecting the combined geometric relationships among different spatial locations in 2D or 3D space instead of the representations of individual spatial feature dimensions. For 2D geometric distance RDM, we computed the 2D Euclidean distance in the 2D space between every pair of stimulus locations as the dissimilarity. Similarly, for 3D geometric distance RDM, we computed the 3D Euclidean distance in the 3D space between every pair of stimulus locations as the dissimilarity.

#### Neural RDMs

EEG electrodes and fMRI voxels each aggregate signals from a mixture of neural sources and spatial locations, making it important to examine both univariate and multivariate dissimilarity measures of representational dissimilarity to consider two types of potential representational formats in the neural recordings. Univariate measures, such as mean amplitude differences, are sensitive to overall or large-scale differences in response strength, while multivariate measures, such as voxel-wise correlation, capture distributed spatial patterns across the scalp (for EEG) or a region of interest (from fMRI). Accordingly, we computed two types of RDMs for each timepoint in the EEG time series and each searchlight unit in the fMRI volume: one based on amplitude differences – the amplitude-based RDM, and one based on correlation distance – the pattern-based RDM.

##### EEG timepoint-by-timepoint RDMs

The EEG neural data consisted of the ERP signal for each of the 64 stimulus locations × 63 electrode channels × 100 timepoints. For the amplitude-based EEG RDMs, we discarded all spatial pattern information. At each timepoint, we computed the mean amplitude across all 63 channels for each stimulus location condition. The dissimilarity between any pair of conditions was then defined as the absolute difference between their mean amplitudes, yielding a 64 × 64 EEG amplitude-based RDM. For the pattern-based EEG RDMs, we discarded absolute activation information by first z-scoring the values at each channel separately across all conditions. At each timepoint, EEG RDMs were computed using a cross-validated, noise-normalized multivariate pattern dissimilarity measure across the 63 channels (Guggenmos et al., 2018). Repetitions for each of the 64 conditions were randomly split into two independent halves, condition-wise mean patterns were computed separately, and dissimilarities were estimated across the two halves after noise normalization using a Ledoit-Wolf shrinkage covariance estimator. This procedure was repeated 50 times with different random splits, and the resulting RDMs were averaged.

##### fMRI searchlight RDMs

The fMRI neural data consisted of BOLD activation beta weights for each of the 64 stimulus locations × 432,216 voxels in the brain. A searchlight analysis was performed using a local searchlight centered on each voxel. Each 3D searchlight consisted of the center voxel and its immediately adjacent neighbors in a 3 × 3 × 3 voxel cube, yielding 27 voxels per searchlight. Searchlights were evaluated at every eligible voxel and were therefore overlapping rather than spatially tiled. We used this relatively local searchlight to preserve spatial specificity when mapping representational structure across cortex.

For the amplitude-based fMRI RDMs, we discarded spatial pattern information within each searchlight unit. For each stimulus location condition, we computed the mean activation (beta value) across all 27 voxels within a given searchlight unit. The dissimilarity between any pair of conditions was defined as the absolute difference between their mean activations, resulting in a 64 × 64 amplitude-based RDM for each searchlight. For the pattern-based fMRI RDMs, we discarded absolute activation information by first z-scoring voxel responses across conditions within each searchlight. Each condition-specific response was treated as a multivoxel pattern, and dissimilarity between pairs of conditions was computed as one minus the Pearson correlation between their voxel-wise activation vectors, yielding a 64 × 64 pattern-based RDM per searchlight unit (across brain space).

##### fMRI ROI-based RDMs

In addition to the searchlight RDMs, we computed a set of RDMs within predefined ROIs (defined above). For each ROI and participant, we extracted the voxel-wise beta weights for all 64 spatial location conditions. For the amplitude-based RDMs, we discarded voxel-level spatial pattern information by computing the mean beta value across all voxels in the ROI for each condition. Dissimilarity between condition pairs was defined as the absolute difference between their mean activation values, resulting in a 64 × 64 amplitude-based RDM per ROI. For the pattern-based RDMs, we first z-scored the voxel-wise beta values across conditions within each ROI to remove mean-level activation differences. Each condition-specific voxel activation pattern was then compared using one minus the Pearson correlation coefficient, yielding a 64 × 64 pattern-based RDM for each ROI.

#### Partial correlations between neural and feature RDMs

To evaluate how human brains represent different spatial features across time (EEG) and brain space (fMRI), we calculated the representational similarity between each neural RDM – temporal EEG RDMs, fMRI searchlight RDMs, or fMRI ROI RDMs – and the ten hypothesis-based feature RDMs described above. To isolate the unique contribution of each spatial feature and mitigate collinearity among feature RDMs, we used rank-based Spearman partial correlation analysis (Dobs et al., 2019; Lu & Golomb, 2024). Spearman correlation was chosen because it is robust to non-linear relationships and differences in scale across RDMs, making it well-suited for comparing representational dissimilarity structures. Specifically, for each 64 × 64 RDM, we extracted the upper triangular values excluding the diagonal (2,016 dissimilarity values) and reshaped them into a 1×2,016 vector. We then computed the partial correlation between a given neural RDM vector and each target feature RDM vector (e.g., the *x* RDM), while statistically controlling for the remaining nine vectors corresponding to the other nine feature RDMs. This procedure removed shared variance with other spatial features and yields a measure of the unique representational similarity for the target feature dimension for each EEG timepoint and fMRI searchlight or ROI.

Similarly, to evaluate whether there is also evidence of processing 2D and 3D location holistically which reflects the combined geometric relationships in 2D or 3D space rather than the individual feature dimensions, we calculated the partial correlation between a given neural RDM vector and each geometric distance RDM vector (2D or 3D geometric distance RDM), while statistically controlling for the alternative geometric distance RDM. Additionally, we also conducted a version of this analysis taking the partial correlation between a given neural RDM vector and each geometric distance RDM vector, while statistically controlling for all feature RDM vectors as well. This procedure removed shared variance with all individual spatial features and the alternative integration representation and yields a measure of the unique representational similarity for the 2D or 3D geometric distance representation regardless of the feature- or coordinate-level encoding format.

As noted above, we constructed two versions of the neural RDMs at each EEG timepoint and fMRI searchlight/ROI – an amplitude-based RDM and a correlation-based RDM. Rather than choose one over the other, since they both capture potentially relevant aspects of the neural signal, we conducted the partial correlation analysis described above for both types of neural RDMs separately and then took the maximum. We defined our robust summary estimate of the final representational similarity for each feature × timepoint (and feature × brain region) as the maximum of these two measures rather than the average because the two measures capture complementary aspects of the neural signal. By taking the maximum, we ensure that if one metric robustly reflects the underlying representation in a given time window or a given brain region – despite potential noise or variations in sensitivity – the overall analysis will not underestimate the representational strength. This conservative approach minimizes the risk of overlooking a genuine representation that may be captured preferentially by one metric over the other. Figures in the main text indicate which type of RDM primarily drove each significant effect, and separated results for the two RDM types are provided in the Supplementary Figures (Figure S8–S13).

#### Statistical Significance: Permutation Tests

To assess the statistical significance of group-level representational similarity for each spatial feature, we performed permutation-based tests. For each permutation, we generated a new random reordering of the 64 stimulus condition labels and then re-conducted the same analysis pipeline described above. In other words, the reordered condition labels were applied identically to both the rows and columns of each neural RDM (EEG temporal RDMs, fMRI searchlight RDMs, or fMRI ROI RDMs), and then we vectorized the upper triangle of the permuted RDM. This procedure preserves the symmetry of the RDM while disrupting the correspondence between neural dissimilarities and the hypothesis-based model RDMs (Nili et al., 2014). We repeated this procedure 1,000 times to generate a null distribution. For each permuted neural RDM, we computed the partial correlation between the permuted neural RDM and each of the ten model RDMs, while controlling for the remaining nine as described above. The resulting partial correlations were then averaged across participants to form a group-level null distribution for each spatial feature. To evaluate significance, we compared the true group-level partial correlation (based on unshuffled neural RDMs) to its corresponding null distribution. To further control for multiple comparisons across hypothesis models, we additionally corrected the significance threshold within each analysis family according to the number of hypothesis-based RDMs tested against each neural RDM. For instance, for the ten feature-level hypothesis-based RDMs, significance was assessed at a Bonferroni-corrected threshold of *p* < 0.05/10. Therefore, a feature was considered significantly represented if the true group-level similarity exceeded the 99.5th percentile of the null distribution (one-sided test, Bonferroni-corrected). For EEG temporal and fMRI searchlight analyses, we further applied cluster-based correction across contiguous timepoints or spatial units to control for multiple comparisons. For ROI-based analyses, no cluster correction was applied due to the absence of spatial or temporal continuity. In addition, Figure S14–S17 show individual representational similarity results and number of significant subjects of each result in time and brain space.

## Data Availability

All data and code are available on GitHub: https://github.com/ZitongLu1996/3D_Visual_Perception.

## Acknowledgements

This work was supported by research grants from the National Institutes of Health (R01-EY025648).

## Supplementary

**Figure S1.**
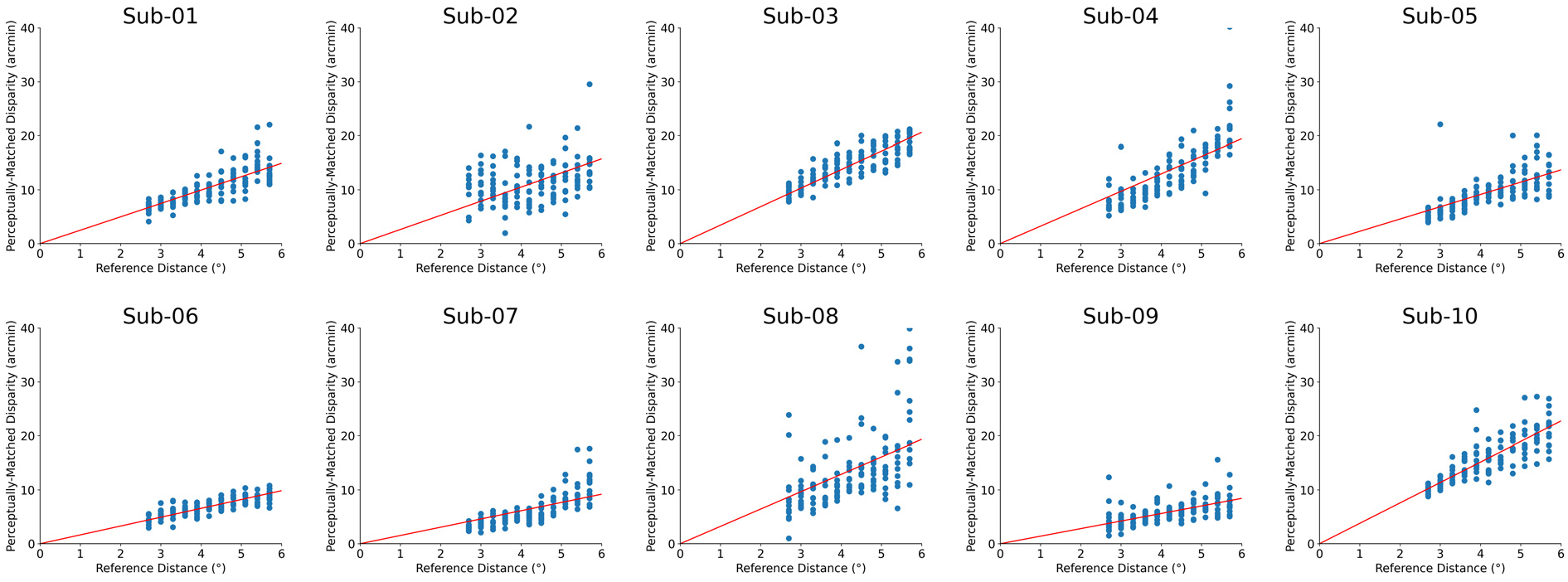
Individual disparity-depth functions derived from the cube adjustment Task. Each panel shows data from one participant (Sub-01 to Sub-10), plotting the relationship between reference distance (in degrees of visual angle) and the perceptual matched binocular disparity (in arcmin) required to perceive a cube with equal 2D and depth dimensions. Each blue dot represents a single trial in which participants adjusted the relative front-back disparity of four RDS patches to form a perceptually cube shape. Red lines indicate the best-fitting linear function (intercept fixed at zero) obtained via least squares regression. These individualized functions were used to convert 3D spatial positions into participant-specific disparity values in the EEG and fMRI main task.

**Figure S2.**
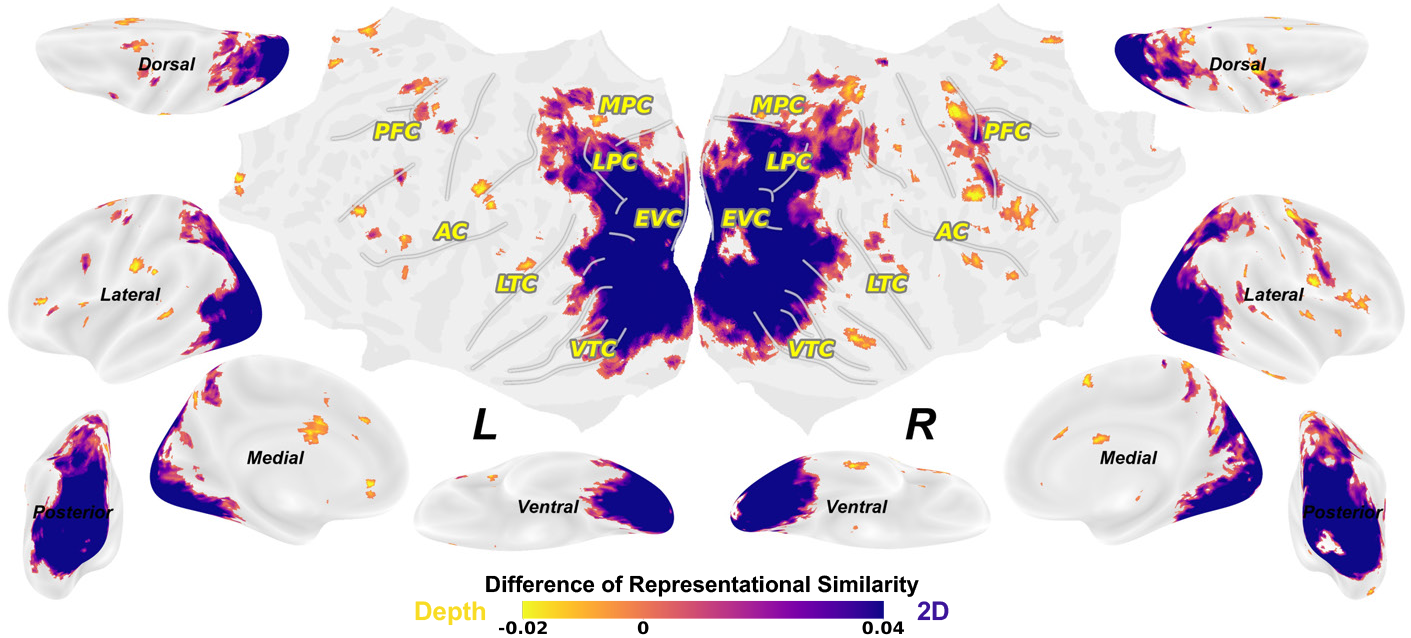
2D minus depth representations from correlation-based RSA results. Additional contrast analysis between 2D (averaging *x* and *y*) and depth (*z*) based on correlation-based RSA results replicated the finding of the spatial transition from 2D to depth along the visual hierarchy in (Finlayson et al., 2017). Colored voxels indicate the searchlight centered on that voxel was significant (permutation tests with cluster-based correction, *p*<.05).

**Figure S3.**
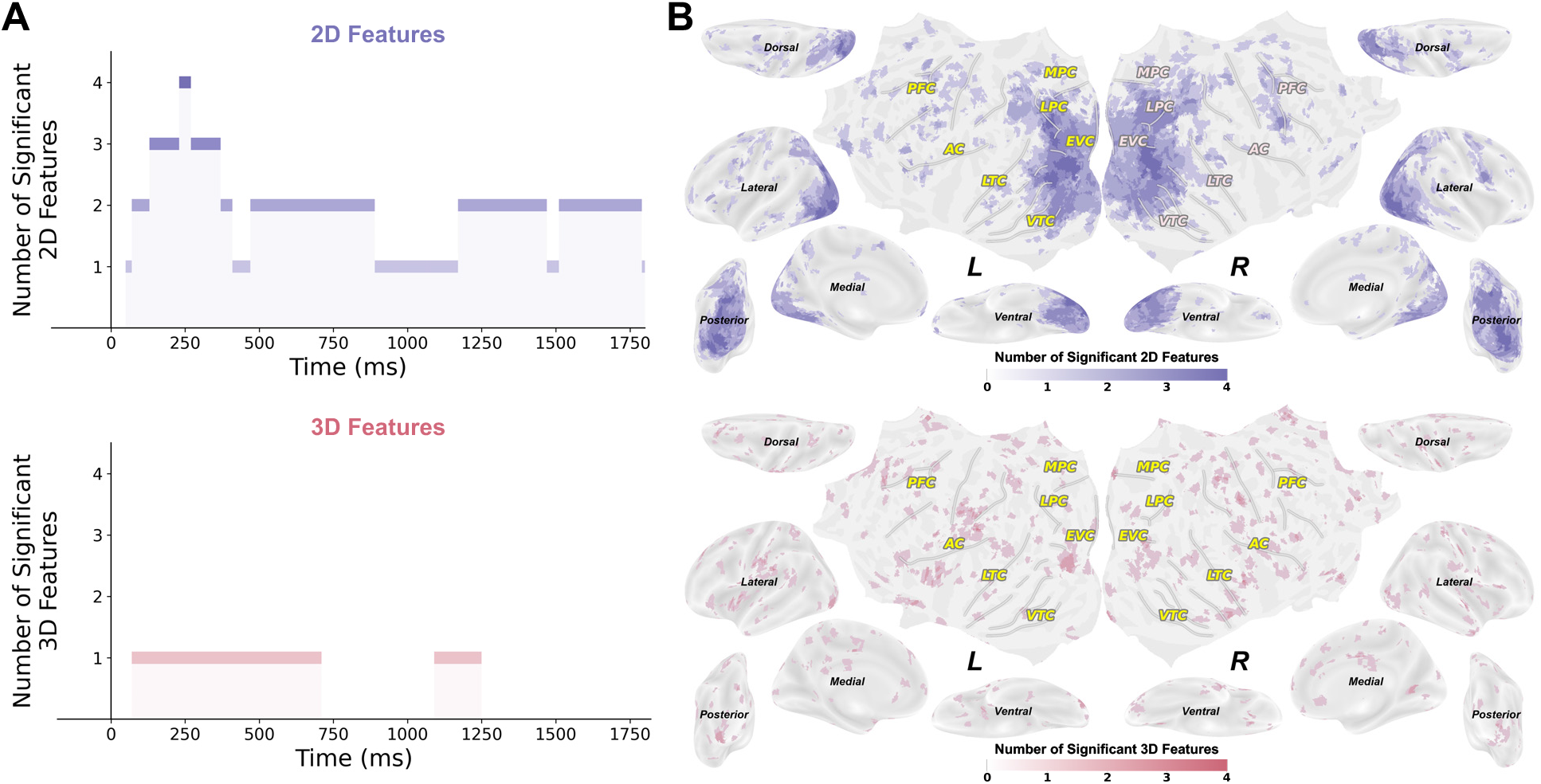
Number of significant 2D or 3D spatial features. (A) Number of significant 2D (upper) or 3D (bottom) spatial features over time from EEG. (B) Number of significant 2D (upper) or 3D (bottom) spatial features across brain space from fMRI.

**Figure S4.**
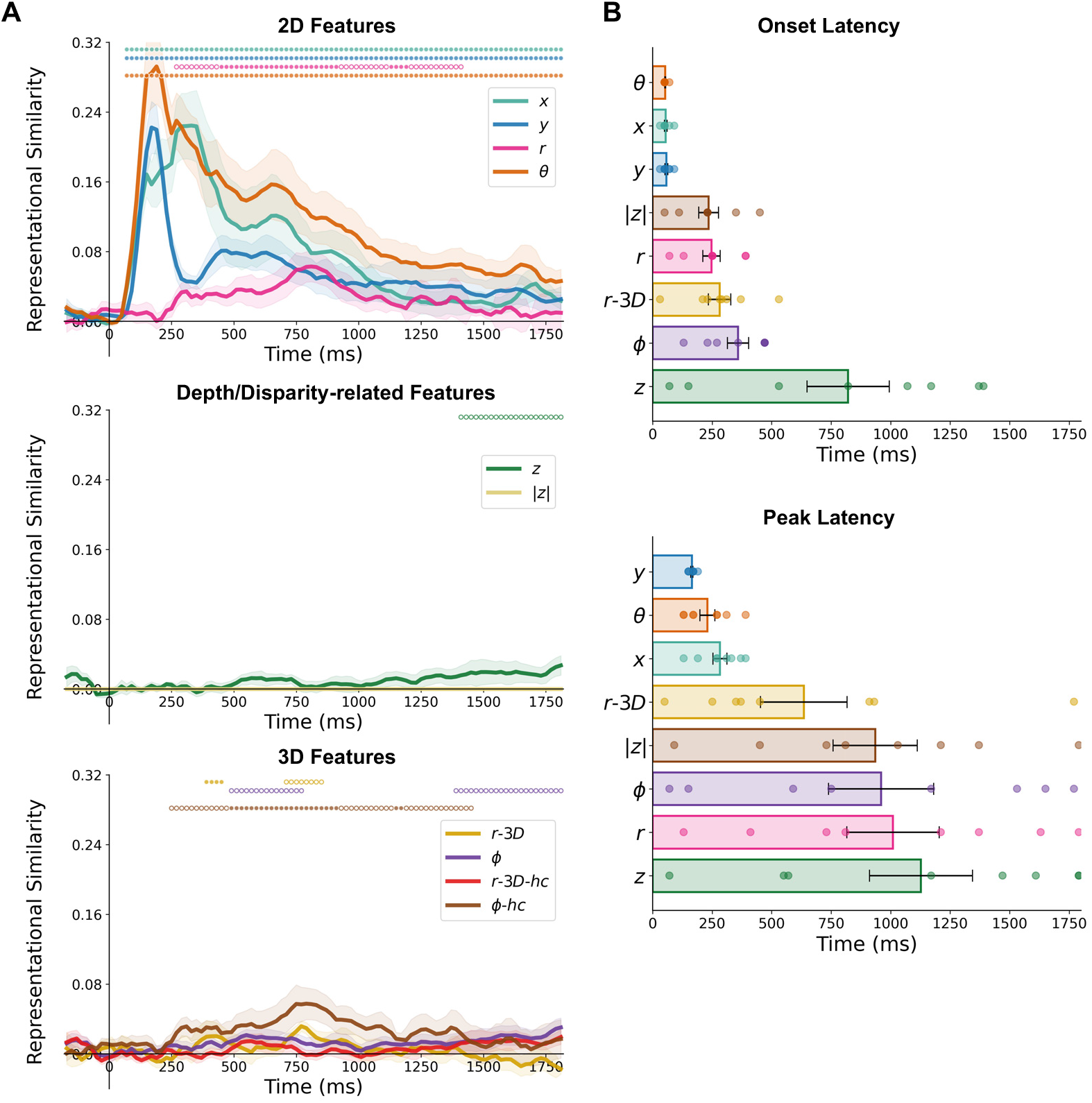
Correlation-based EEG RSA results of spatial features without controlling for shared variances among model RDMs. (A) Time-resolved representational similarity (Spearman correlation) between EEG temporal RDMs and the ten hypothesis-based spatial feature RDMs, including 2D features (*x*, *y*, *r*, *θ*), depth/disparity-related features (*z* and |*z*|), and 3D features (*r-3D*, *ϕ*, *r-3D-hc*, *ϕ-hc*). Shaded areas indicate ±1 SEM across participants. Open circles indicate significant time points primarily driven by amplitude-based RDMs, whereas filled circles indicate significant time points primarily driven by pattern-based RDMs (permutation tests with correction for the number of hypothesis-based RDMs, followed by cluster-based correction, *p*<.05). (B) Onset and peak latencies of EEG representational similarity effects for spatial features that showed significant time points. Features without significant EEG effects (|*z*| and *r-3D-hc*) were not included in the latency summary. Dots denote individual participants; error bars indicate SEM. **This figure corresponds to Figure 3 in the main text but uses pure Spearman correlation without controlling for shared variances among model RDMs.**

**Figure S5.**
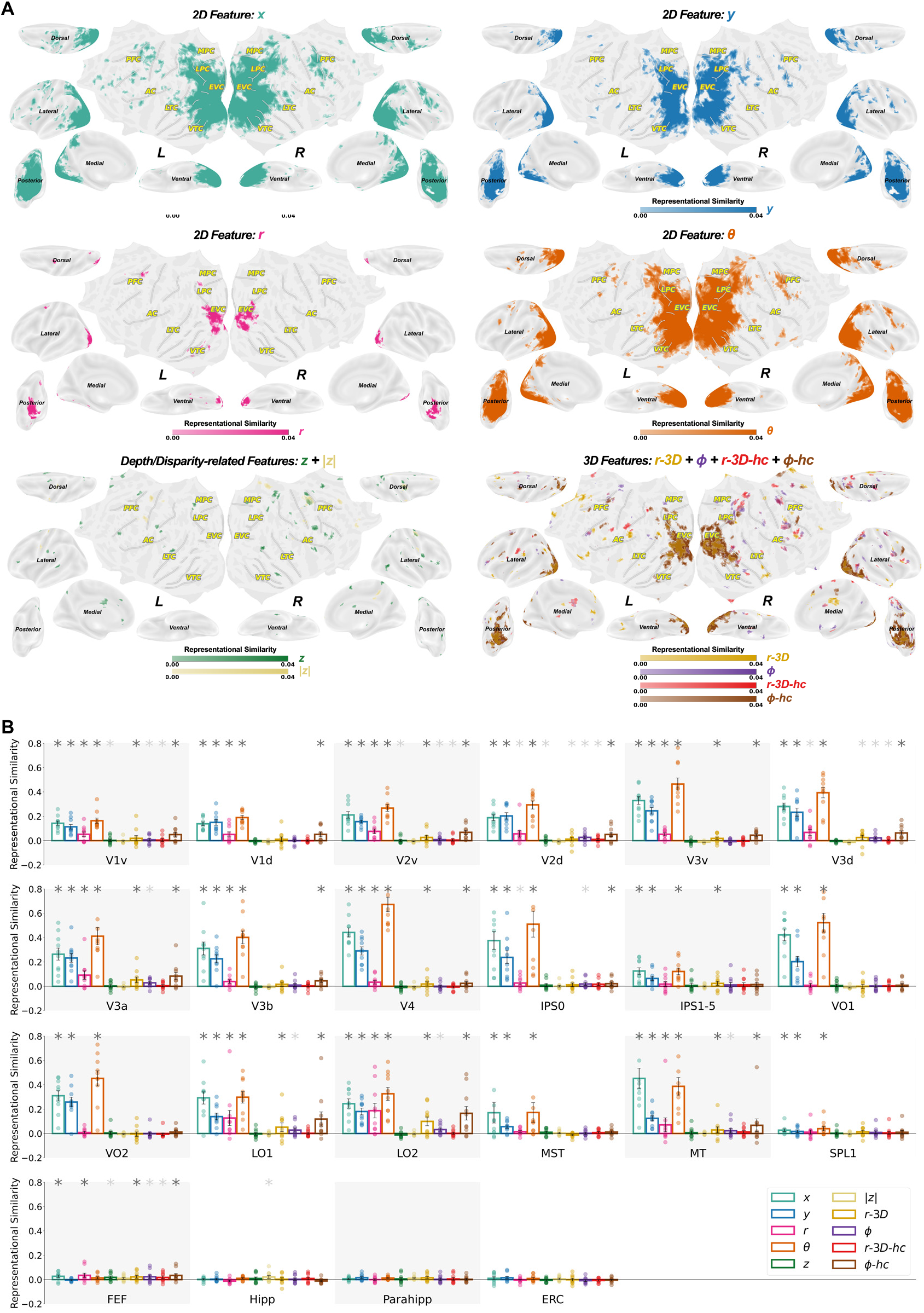
Correlation-based fMRI RSA results of individual spatial feature representations without controlling for shared variances among model RDMs. (A) fMRI searchlight representational similarity (Spearman correlation) results for spatial features displayed on inflated cortical surfaces. The four 2D feature maps (*x*, *y*, *r*, *θ*) are shown separately, whereas depth/disparity-related features (*z* and |*z*|), and 3D features (*r-3D*, *ϕ*, *r-3D-hc*, *ϕ-hc*) are summarized as two combined presence maps because their individual maps were more sparse and heterogeneous. Colored voxels indicate the searchlight centered on that voxel was significant (permutation tests with correction for the number of hypothesis-based RDMs, followed by cluster-based correction, *p*<.05). (B) ROI-based RSA results showing group-level representational similarity (Spearman correlation) across 25 predefined ROIs. Light-gray asterisks indicate significant effects primarily driven by amplitude-based RDMs, whereas dark-gray asterisks indicate significant effects primarily driven by pattern-based RDMs (permutation tests with correction for the number of hypothesis-based RDMs, *p*<.05). Dots denote individual participants. Error bars indicate SEM. **This figure corresponds to Figure 4 in the main text but uses pure Spearman correlation without controlling for shared variances among model RDMs.**

**Figure S6.**
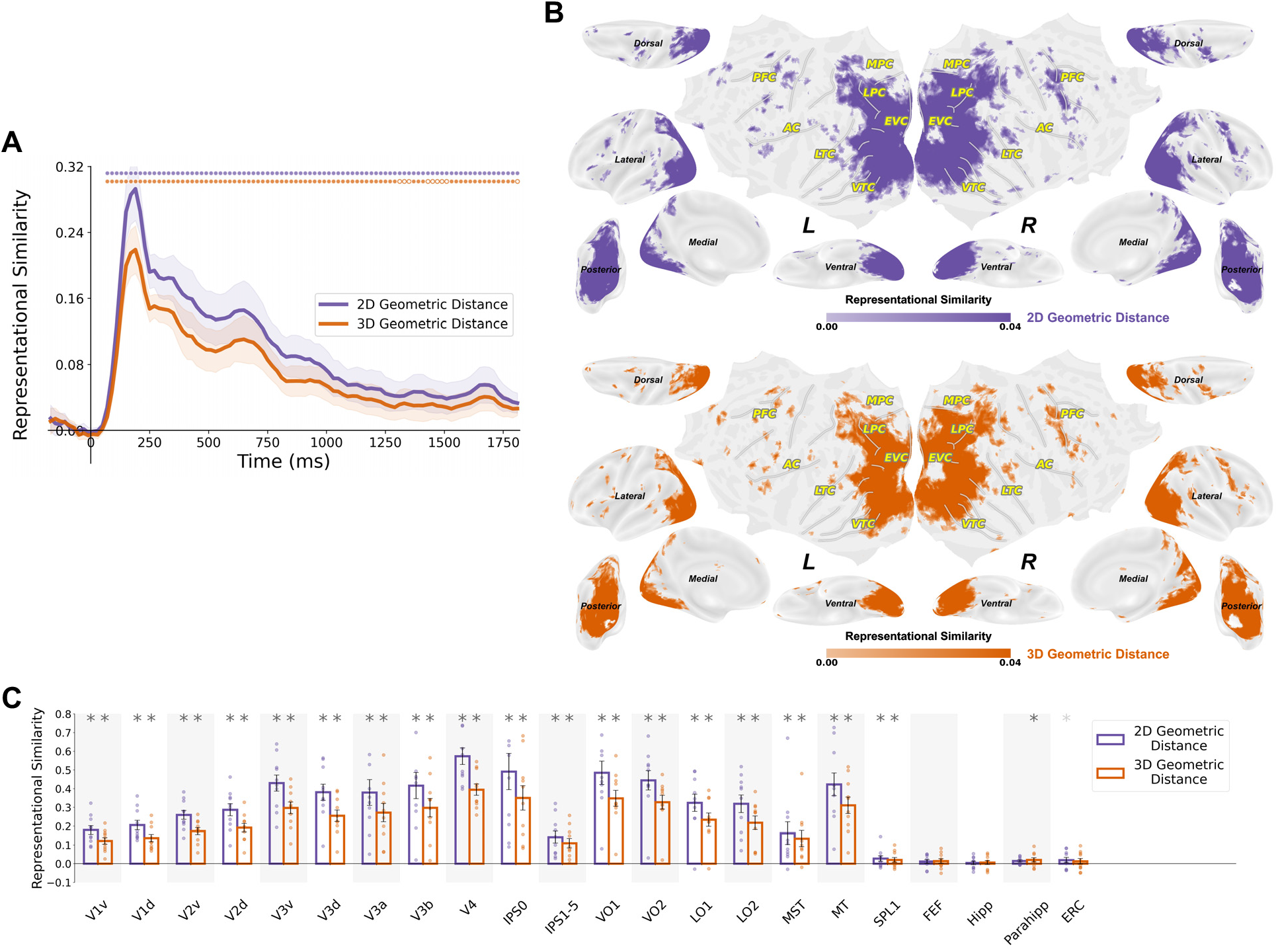
Correlation-based RSA results of 2D and 3D geometric distance representations without controlling for shared variances among model RDMs. (A) Time-resolved representational similarity (Spearman correlation) between EEG temporal RDMs and the two geometric distance RDMs. Shaded areas indicate ±1 SEM across participants. Open circles indicate significant time points primarily driven by amplitude-based RDMs, whereas filled circles indicate significant time points primarily driven by pattern-based RDMs (permutation tests with correction for the number of hypothesis-based RDMs, followed by cluster-based correction, *p*<.05). (B) fMRI searchlight representational similarity (Spearman correlation) between fMRI temporal RDMs and the two geometric distance RDMs. fMRI searchlight representational similarity maps displayed on inflated cortical surfaces. Colored voxels indicate the searchlight centered on that voxel was significant (permutation tests with correction for the number of hypothesis-based RDMs, followed by cluster-based correction, *p*<.05). (C) ROI-based RSA results showing group-level representational similarity (Spearman correlation) of 2D and 3D geometric distances across 25 predefined ROIs. Light-gray asterisks indicate significant effects primarily driven by amplitude-based RDMs, whereas dark-gray asterisks indicate significant effects primarily driven by pattern-based RDMs (permutation tests with correction for the number of hypothesis-based RDMs, *p*<.05). Dots denote individual participants. Error bars indicate SEM. **This figure corresponds to Figure 5 in the main text but uses pure Spearman correlation without controlling for shared variances among model RDMs.**

**Figure S7.**
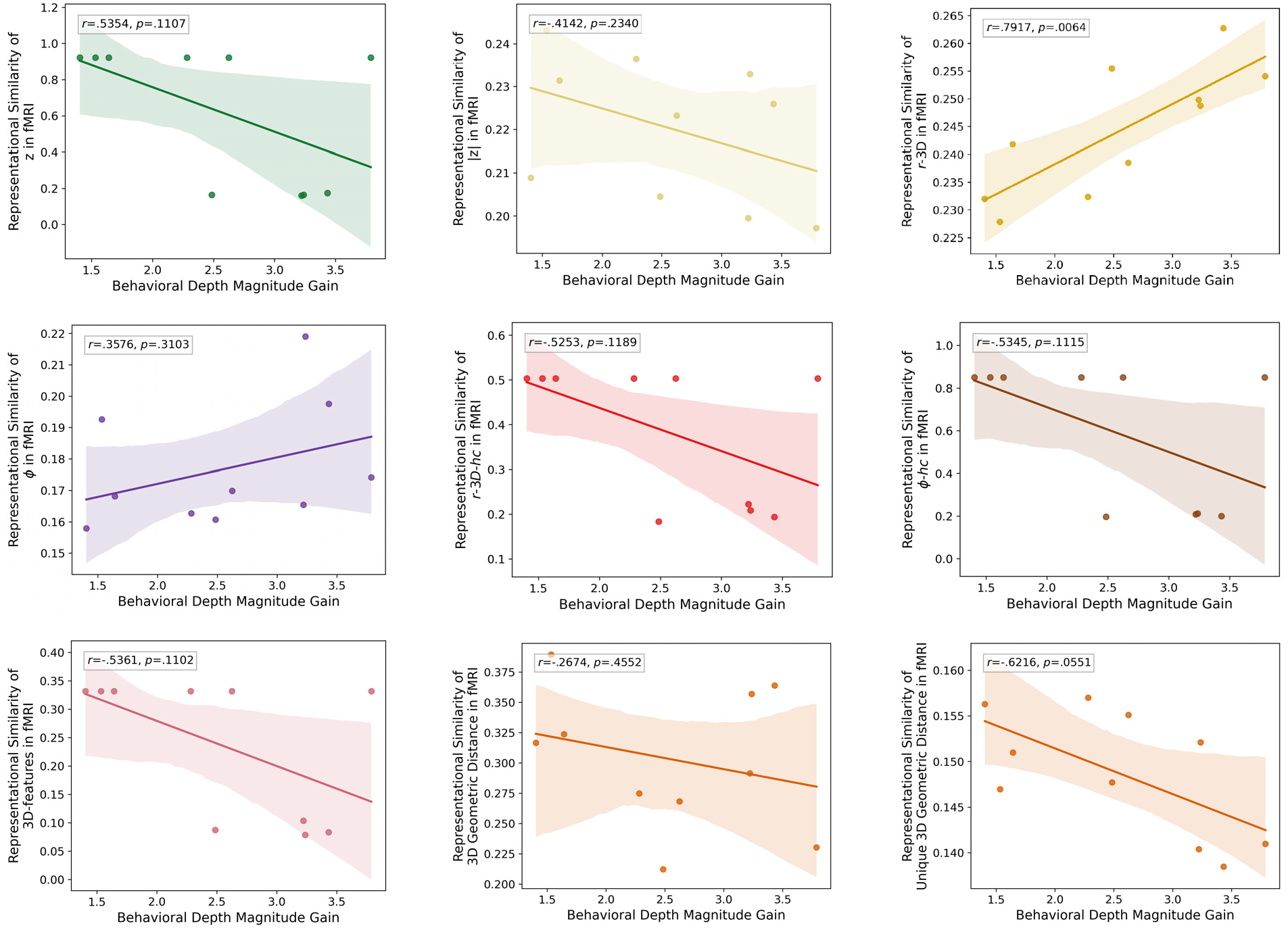
Correlation between behavioral depth magnitude gain and neural encoding in fMRI. Each dot represents one participant. Shaded area indicates 95% confidence interval for the fitted regression line. **This figure corresponds to Figure 6 in the main text but plotted for all depth- and 3D-related RSA results.**

**Figure S8.**
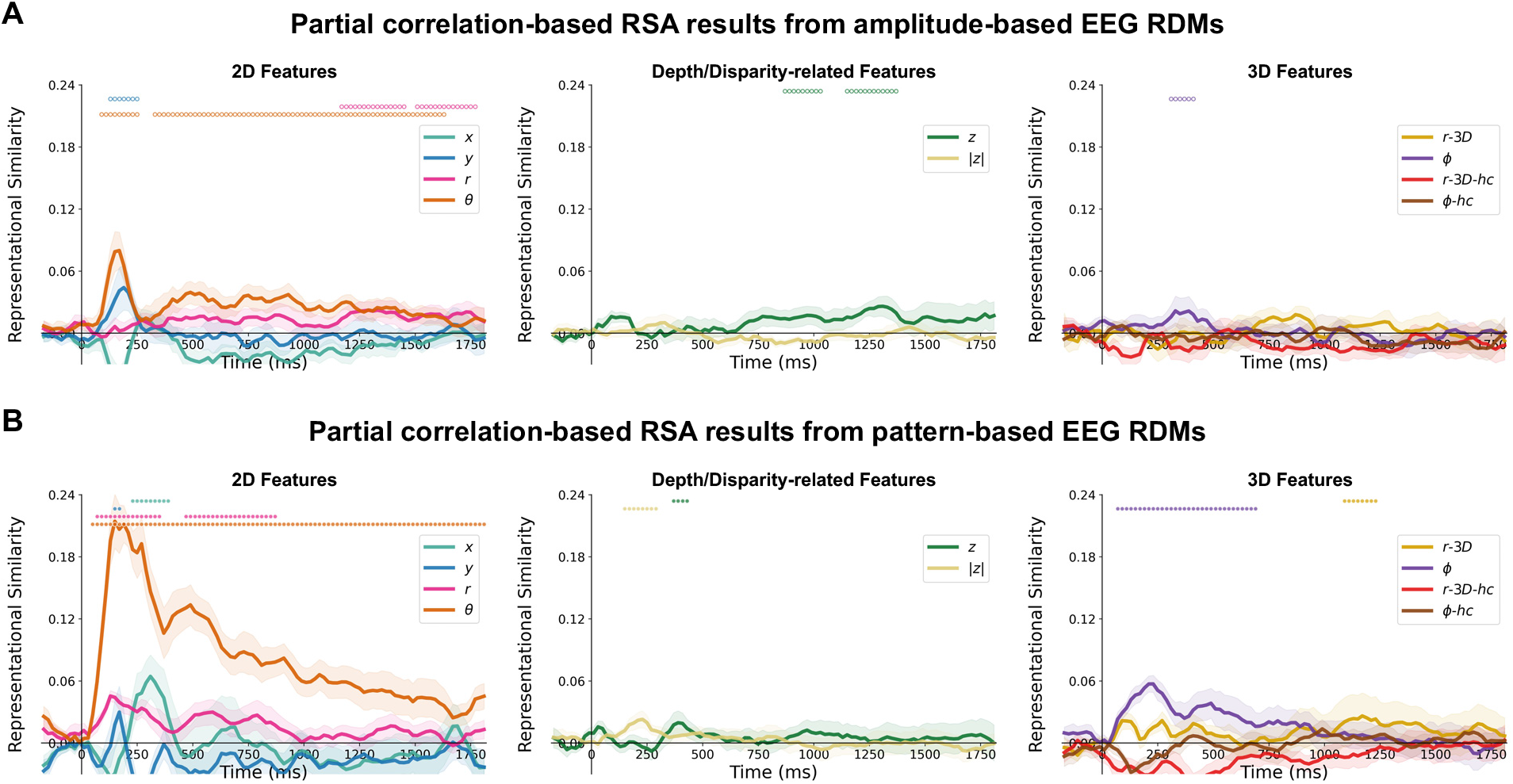
Partial correlation-based EEG RSA results of spatial features from amplitude- and pattern-based EEG RDMs. Time-resolved representational similarity (partial Spearman correlation) results between EEG temporal (A) amplitude- or (B) pattern-based RDMs and ten hypothesis-based feature RDMs, grouped by 2D (*x*, *y*, *r*, *θ*), depth/disparity-related (*z* and |*z*|), and 3D (*r-3D-hc*, *θ*, *Φ-hc*) features. Shaded areas indicate ±1 SEM across participants. Open circles indicate significant time points primarily driven by amplitude-based RDMs, whereas filled circles indicate significant time points primarily driven by pattern-based RDMs (permutation tests with correction for the number of hypothesis-based RDMs, followed by cluster-based correction, *p*<.05). **This figure corresponds to Figure 3A in the main text but plotted separately for amplitude- and pattern-based results.**

**Figure S9.**
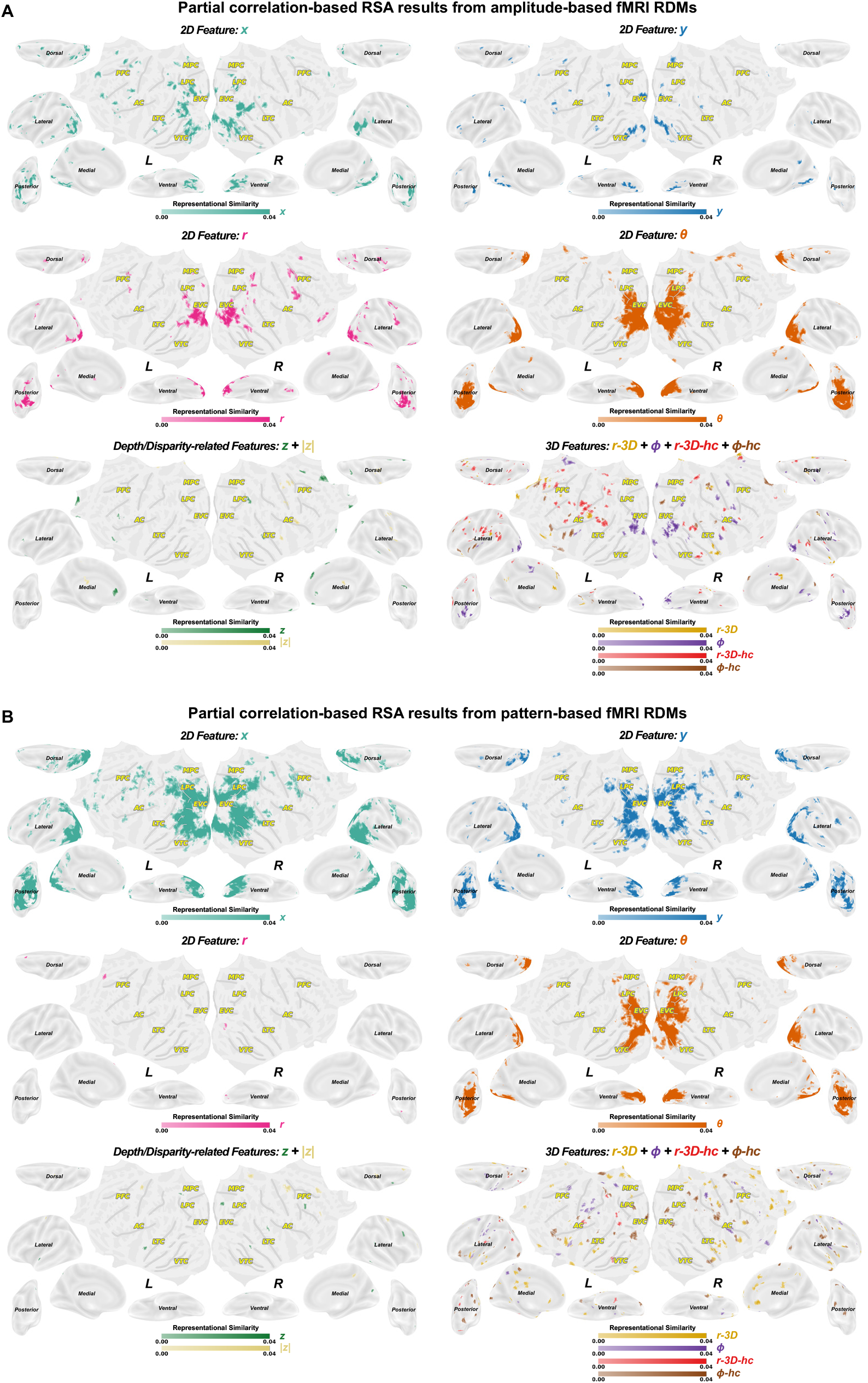
Partial correlation-based fMRI RSA results of individual spatial features from amplitude- and pattern-based fMRI RDMs. Searchlight representational similarity (partial Spearman correlation) results between fMRI searchlight (A) amplitude- or (B) pattern-based RDMs and ten hypothesis-based spatial feature RDMs (permutation tests with correction for the number of hypothesis-based RDMs, followed by cluster-based correction, *p*<.05). **This figure corresponds to Figure 4A in the main text but plotted separately for amplitude- and pattern-based results.**

**Figure S10.**
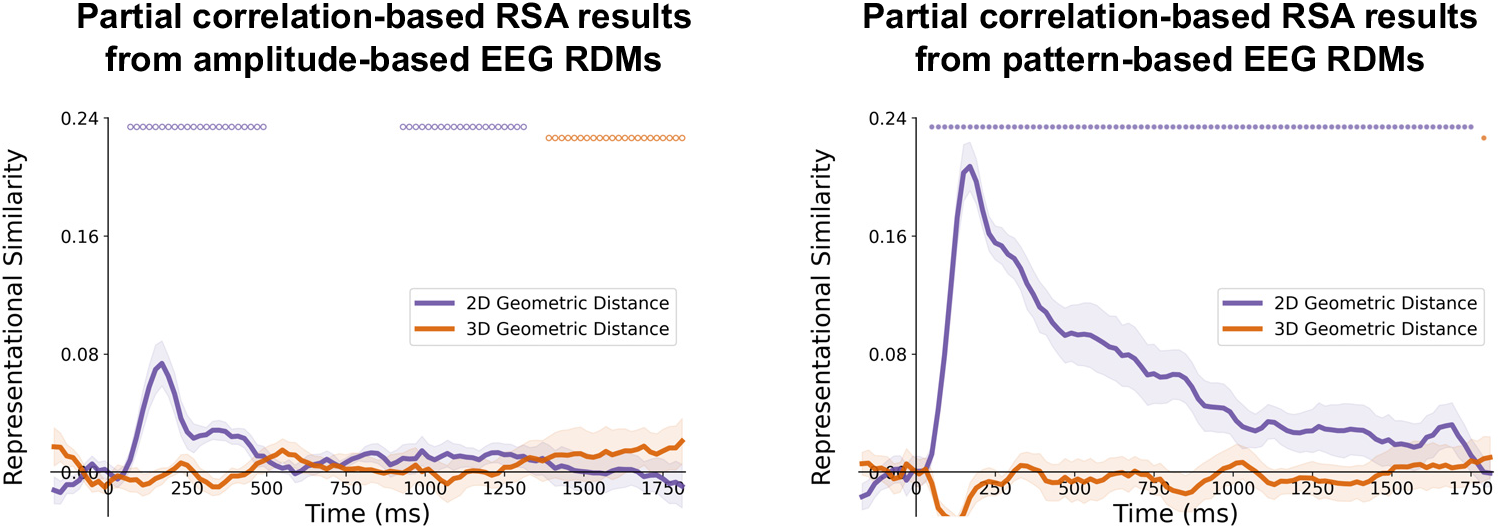
Partial correlation-based EEG RSA results of geometric representation from amplitude- and pattern-based EEG RDMs. Time-resolved representational similarity (partial Spearman correlation) results between EEG temporal amplitude- or correlation-based RDMs and two geometric distance RDMs. Shaded areas indicate ±1 SEM across participants. Open circles indicate significant time points primarily driven by amplitude-based RDMs, whereas filled circles indicate significant time points primarily driven by pattern-based RDMs (permutation tests with correction for the number of hypothesis-based RDMs, followed by cluster-based correction, *p*<.05). **This figure corresponds to Figure 5A in the main text but plotted separately for amplitude- and pattern-based results.**

**Figure S11.**
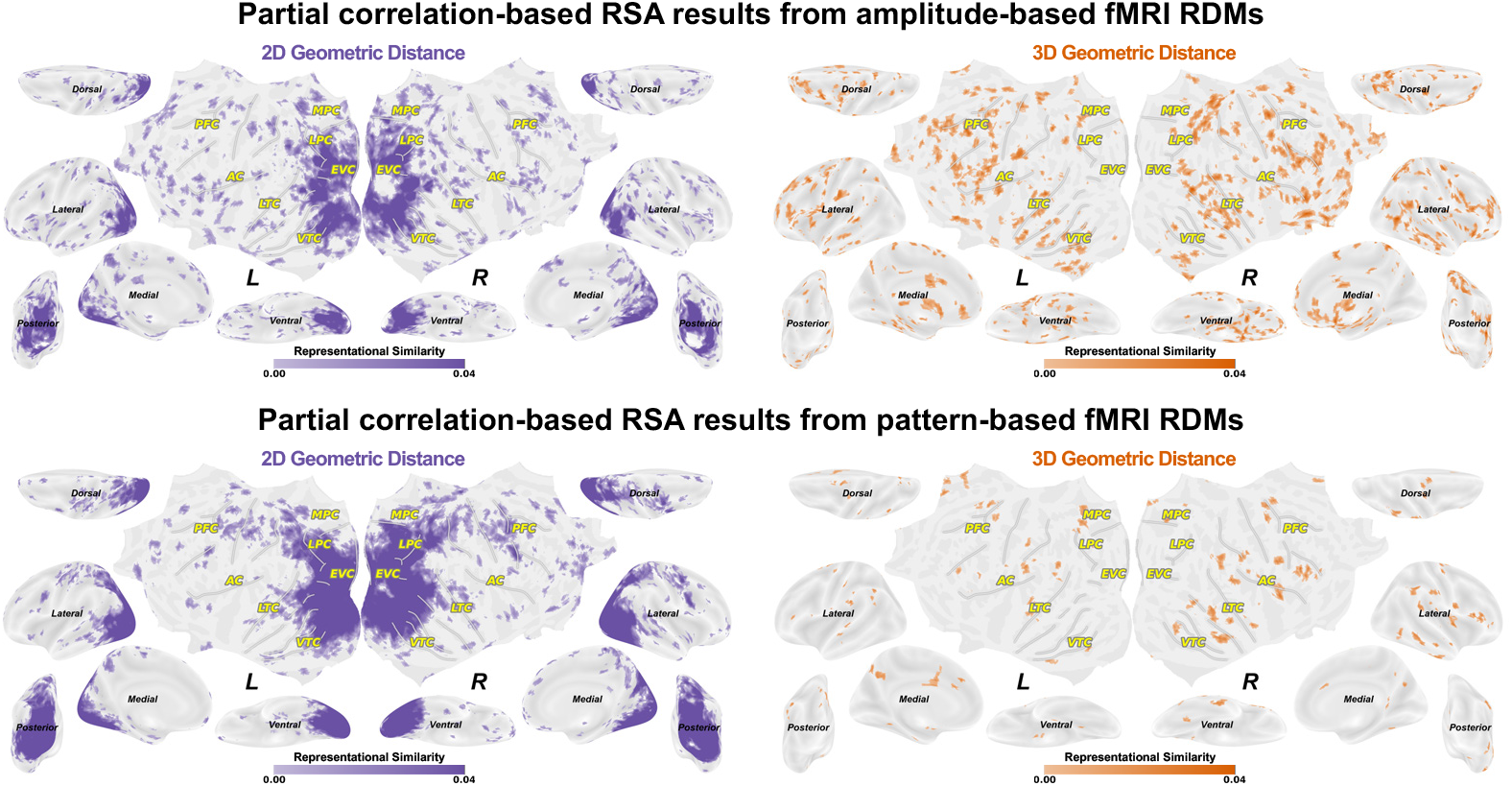
Partial correlation-based fMRI RSA results of geometric distance representation from amplitude- and pattern-based fMRI RDMs. Searchlight representational similarity (partial Spearman correlation) results between fMRI searchlight amplitude- or pattern-based RDMs and two hypothesis-based geometric distance RDMs (permutation tests with correction for the number of hypothesis-based RDMs, followed by cluster-based correction, *p*<.05). **This figure corresponds to Figure 5B in the main text but plotted separately for amplitude- and pattern-based results.**

**Figure S12.**
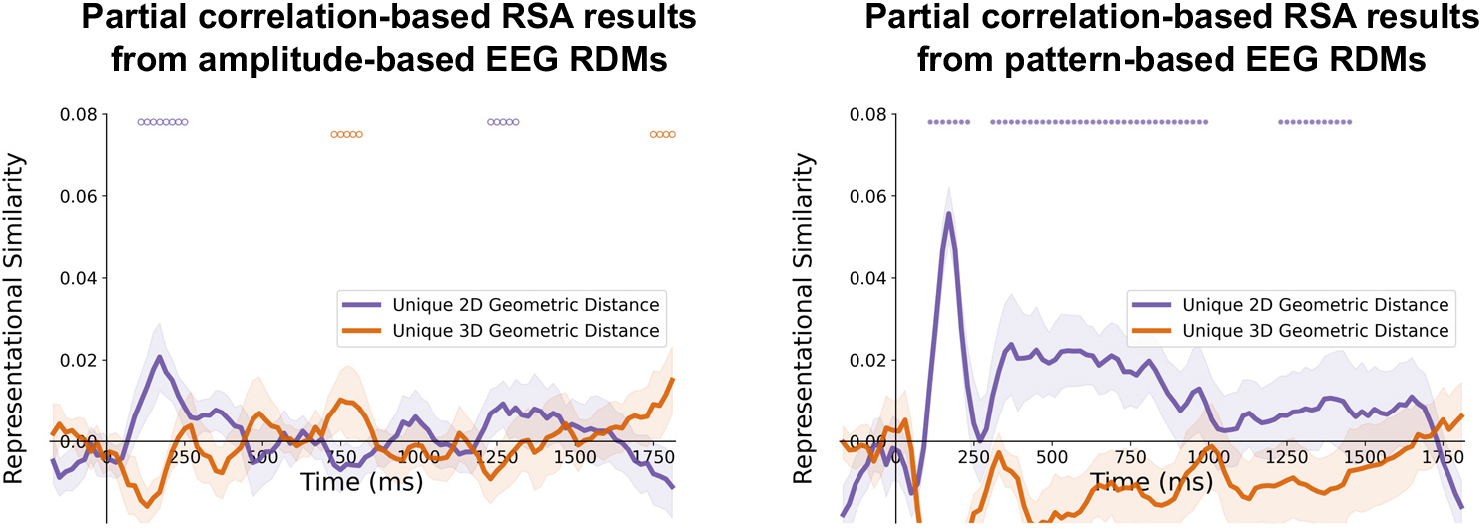
Partial correlation-based EEG RSA results of unique geometric representation from amplitude- and pattern-based EEG RDMs. Time-resolved representational similarity (partial Spearman correlation, also controlling ten feature RDMs) results between EEG temporal amplitude- or correlation-based RDMs and two geometric distance RDMs. Shaded areas indicate ±1 SEM across participants. Open circles indicate significant time points primarily driven by amplitude-based RDMs, whereas filled circles indicate significant time points primarily driven by pattern-based RDMs (permutation tests with correction for the number of hypothesis-based RDMs, followed by cluster-based correction, *p*<.05). **This figure corresponds to Figure 5D in the main text but plotted separately for amplitude- and pattern-based results.**

**Figure S13.**
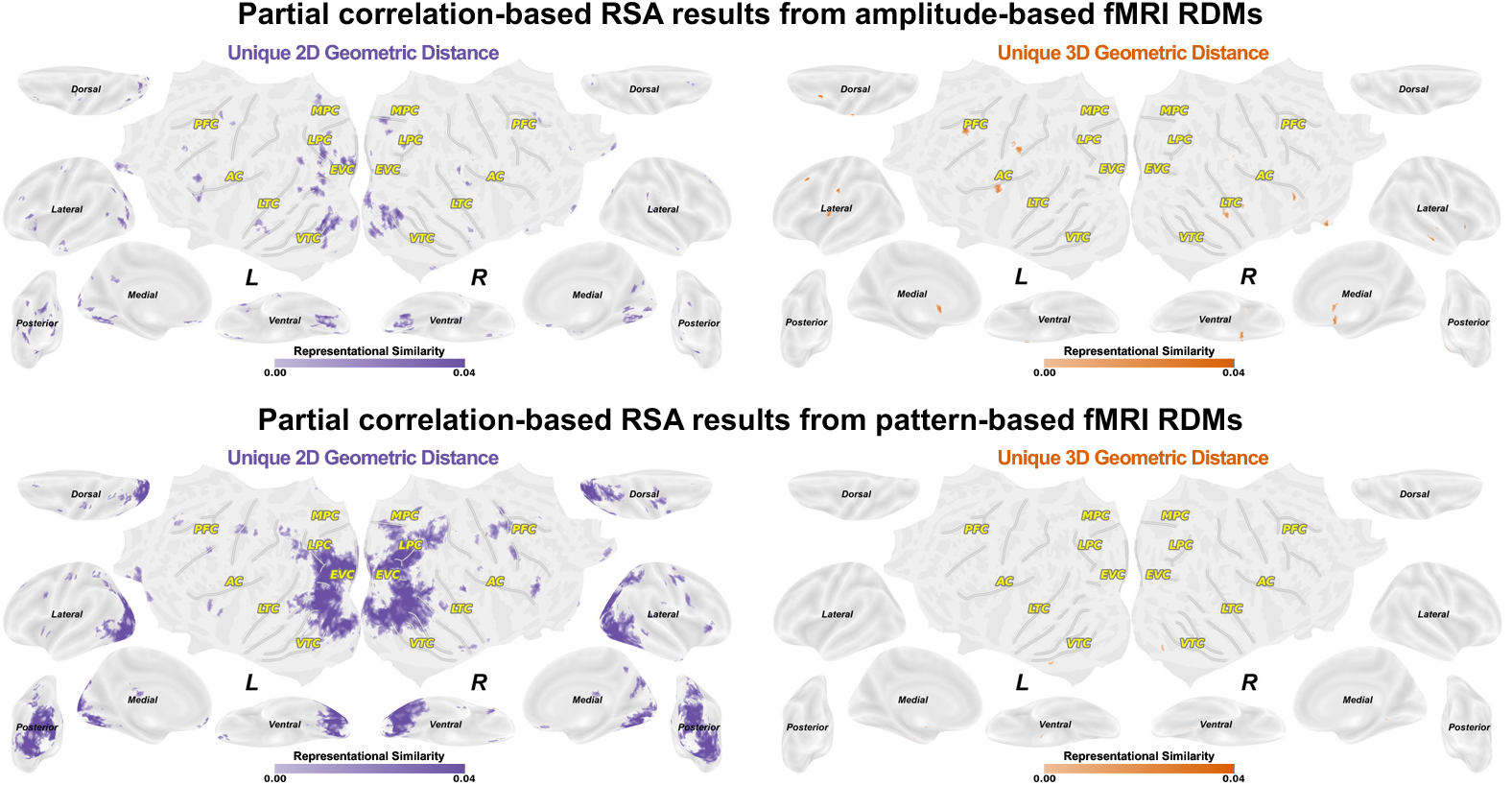
Partial correlation-based fMRI RSA results of unique geometric distance representation from amplitude- and pattern-based fMRI RDMs. Searchlight representational similarity (partial Spearman correlation, also controlling nine feature RDMs) results between fMRI searchlight amplitude- or pattern-based RDMs and two hypothesis-based geometric distance RDMs (permutation tests with correction for the number of hypothesis-based RDMs, followed by cluster-based correction, *p*<.05). **This figure corresponds to Figure 5E in the main text but plotted separately for amplitude- and pattern-based results.**

**Figure S14.**
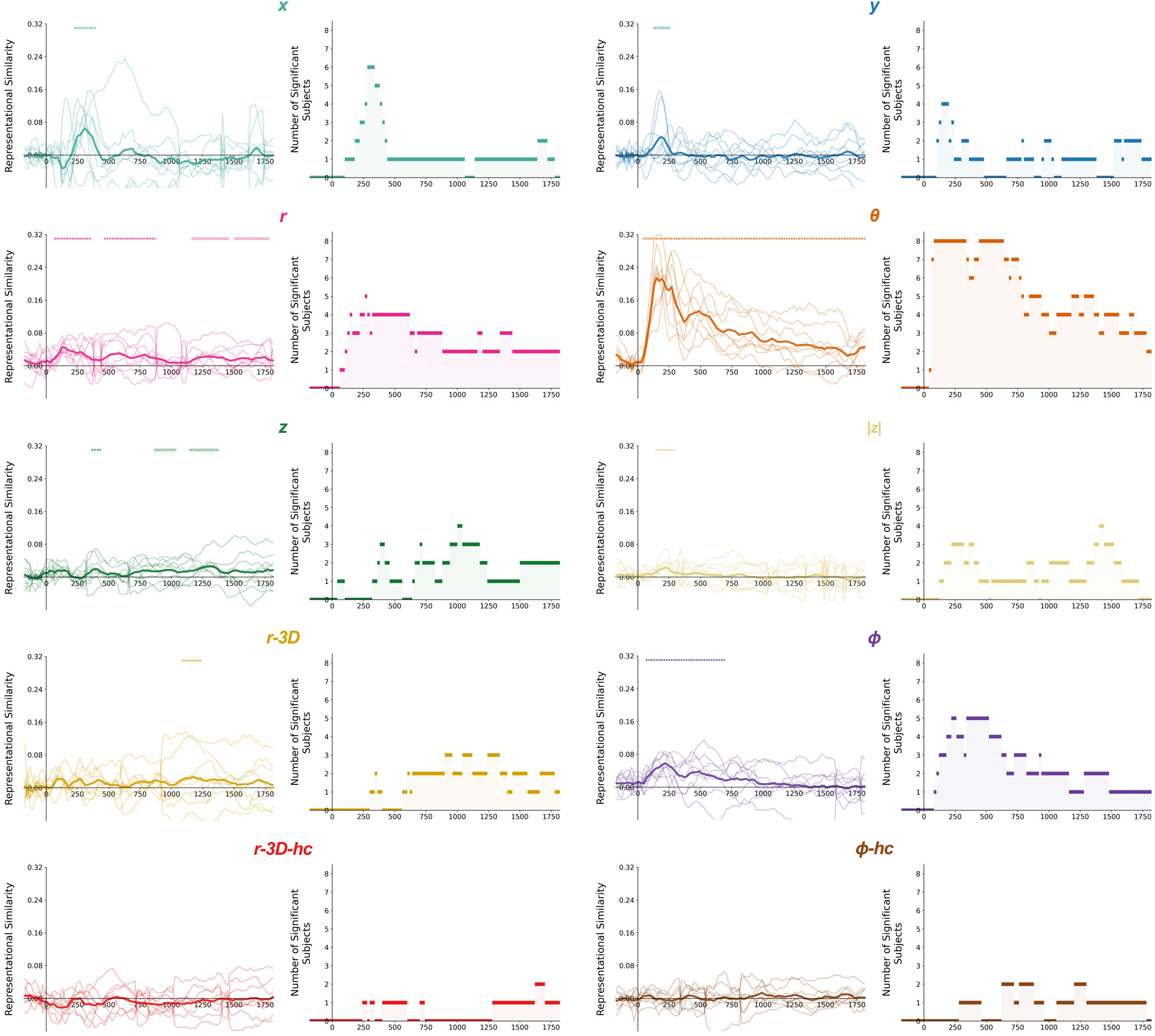
Number of significant subjects of EEG-based partial RSA results of individual spatial features. Each light and thin line correspond to a single participant. The dark and think line correspond to the averaged result across eight participants. Open circles indicate significant time points primarily driven by amplitude-based RDMs, whereas filled circles indicate significant time points primarily driven by pattern-based RDMs (for each subject’s results: permutation tests with correction for the number of hypothesis-based RDMs, followed by cluster-based correction, *p*<.05). **This figure corresponds to Figure 3A in the main text but plotted for number of significant subjects.**

**Figure S15.**
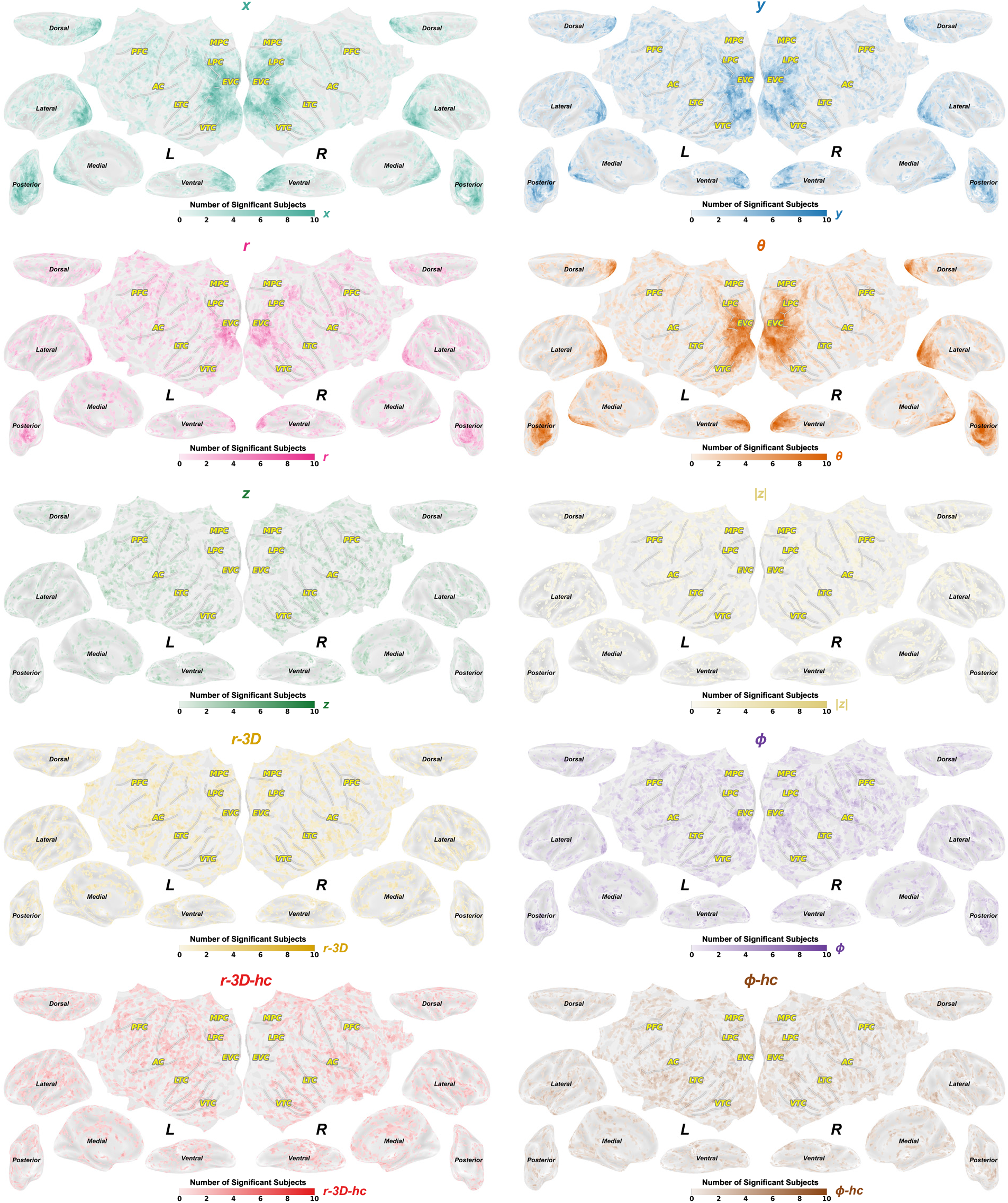
Number of significant subjects of fMRI-based partial RSA results of individual spatial features. (for each subject’s results: permutation tests with correction for the number of hypothesis-based RDMs, followed by cluster-based correction, *p*<.05). **This figure corresponds to Figure 4A in the main text but plotted for number of significant subjects.**

**Figure S16.**
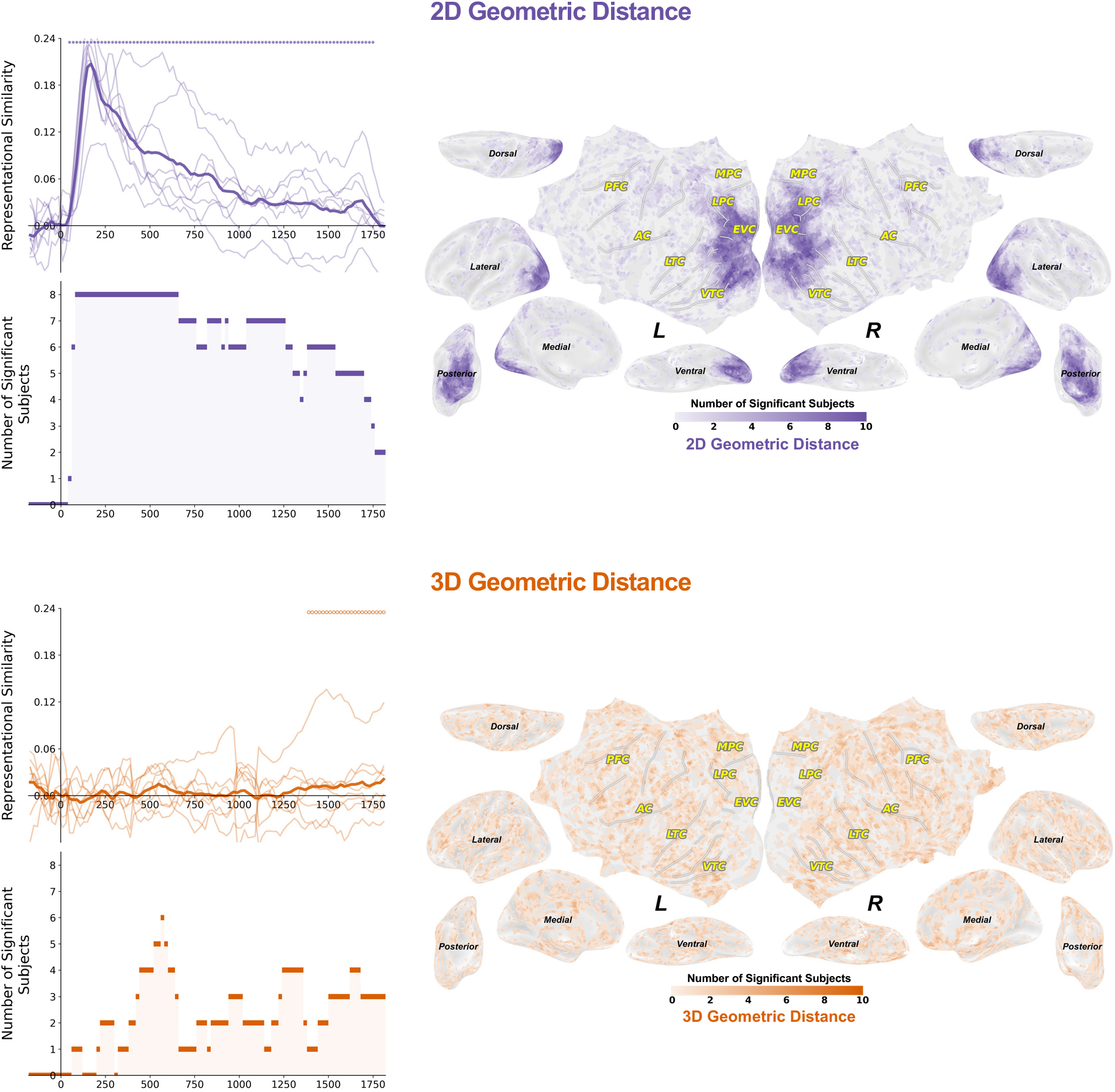
Number of significant subjects of partial RSA results of geometric distance representations. Each light and thin line correspond to a single participant. The dark and think line correspond to the averaged result across eight (EEG) or ten (fMRI) participants. Open circles indicate significant time points primarily driven by amplitude-based RDMs, whereas filled circles indicate significant time points primarily driven by pattern-based RDMs (for each subject’s results: permutation tests with correction for the number of hypothesis-based RDMs, followed by cluster-based correction, *p*<.05). **This figure corresponds to Figure 5A-B in the main text but plotted for number of significant subjects.**

**Figure S17.**
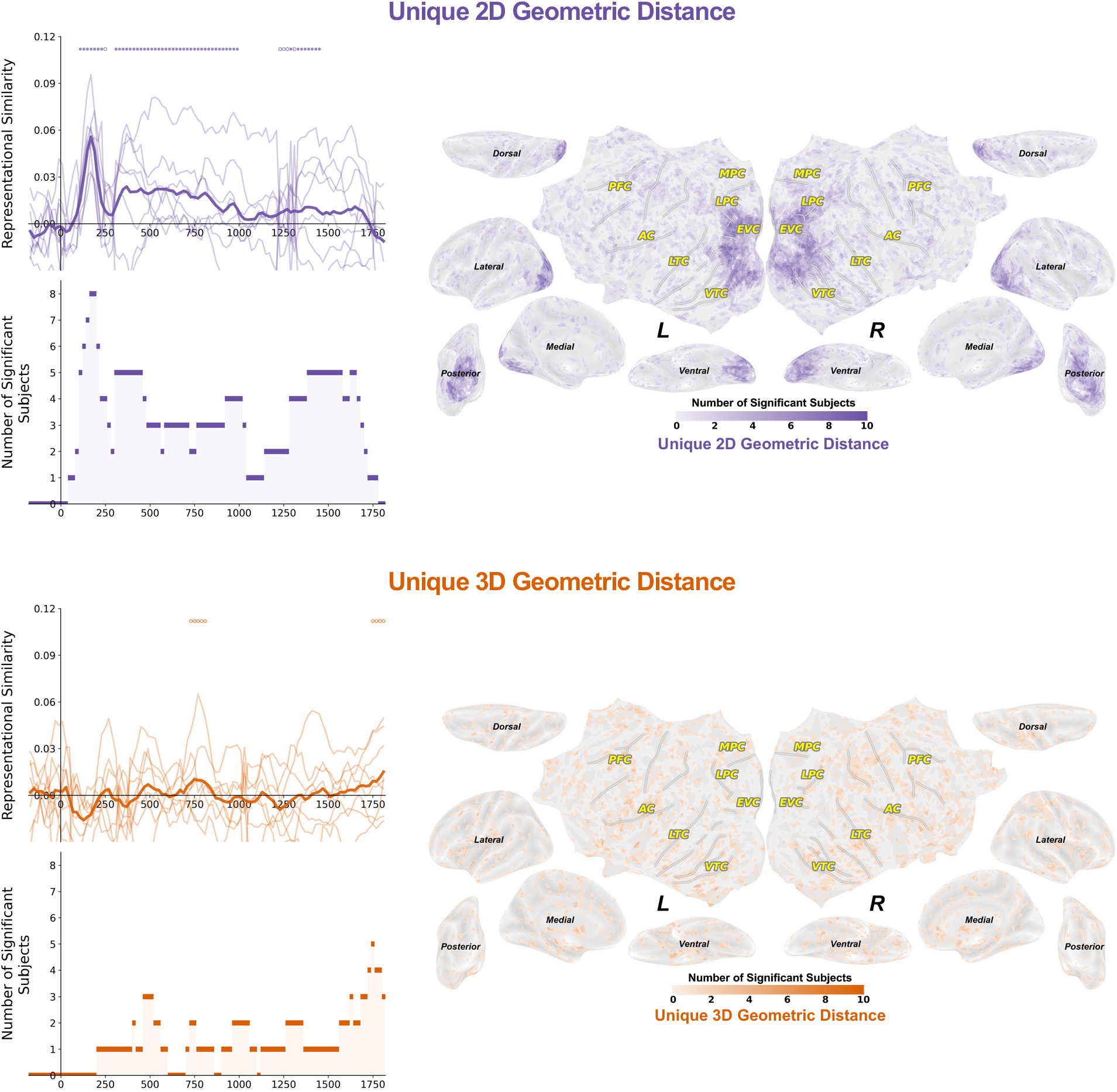
Number of significant subjects of partial RSA results of unique geometric distance representations. Each light and thin line correspond to a single participant. The dark and think line correspond to the averaged result across eight (EEG) or ten (fMRI) participants. Open circles indicate significant time points primarily driven by amplitude-based RDMs, whereas filled circles indicate significant time points primarily driven by pattern-based RDMs (for each subject’s results: permutation tests with correction for the number of hypothesis-based *RDMs*, followed by cluster-based correction, *p*<.05). **This figure corresponds to Figure 5D-E in the main text but plotted for number of significant subjects.**

(See the video via the link: https://github.com/ZitongLu1996/3D_Visual_Perception/blob/master/video.mp4)

**Video S1. Visualization of combined EEG and fMRI RSA results of spatial feature representations.** In this video, if the EEG RSA result of a certain spatial feature is significant, we highlighted the corresponding significant fMRI RSA result of the feature on inflated cortical surfaces (based on Figure 3A and 4A).

